# TGF-β signaling promotes the expression of proviral circular RNA ciTRAN in HIV-1 infection

**DOI:** 10.1101/2024.08.28.610092

**Authors:** Aditi Choudhary, Katyayani Mallick, Rishikesh Dalavi, Ajit Chande

## Abstract

Circular RNA (circRNA) expression is widespread in immune cells infected by HIV-1, but the crosstalk between circRNA expression and various cellular signaling pathways remains unclear. We report that HIV-1 Vpr triggers TGF-β signaling, which we linked to the increased expression of ciTRAN, a proviral circRNA encoded by *SMARCA5*. Consistent with this finding, we observed that the essential intracellular TGF-β receptor signaling component SMAD2/3 was recruited to the *SMARCA5* promoter in a Vpr-dependent manner. *SMARCA5* promoter analysis and functional assays further revealed that the SAMD2/3 binding motif is crucial for ciTRAN upregulation. In response to treatment with DNA-damaging agents or the exogenous addition of recombinant TGF-β, the TGF-β-SMAD axis upregulated the expression of ciTRAN as well as the parental *SMARCA5* mRNA. Finally, pharmacological targeting of TGF-β signaling or genetic ablation of *TGFBR1* can reduce the ability of HIV-1 Vpr to induce the expression of ciTRAN and viral genes. These results offer crucial mechanistic insights into the regulation of ciTRAN expression by TGF-β signaling and suggest a strategy to inhibit circRNA expression during infection by small molecules.

## Introduction

Circular RNAs (circRNAs) are an important subclass of RNAs that are widespread in metazoans(Jeck et al., 2013; Salzman et al., 2012; P. L. Wang et al., 2014). In contrast to linear RNAs, which are produced by canonical splicing, circRNAs are generated by backsplicing, wherein a 5’ splice donor links to a 3’ splice site upstream, generating a circular species of RNA resistant to exonuclease-mediated decay(Dong et al., 2016; Starke et al., 2015; X. Zhang et al., 2014). CircRNAs are evolutionarily conserved and developmentally regulated. Moreover, circRNAs display cell-specific expression patterns, are intrinsically stable and can regulate various cellular processes(Chen, 2020; Lasda & Parker, 2014). In addition to the early studies revealing microRNA sponging as a primary function(Hansen et al., 2013; Memczak et al., 2013) of circRNAs, recent studies reveal many other roles, such as regulators of gene expression, as ribosome templates in various pathophysiological conditions and, more recently, as modulators of viral infection(Choudhary et al., 2021).

The influence of circRNAs derived from both hosts and viruses on infection outcomes are recently reported (Edgar et al., 2022; Lee et al., 2023; Tagawa et al., 2018; Ungerleider et al., 2019; W. Wang et al., 2023). Retroviruses are no exception, and the first report on HIV-1 coopting a circRNA to promote its transcription highlighted the importance of circRNA in retroviral pathogenesis(Bhardwaj et al., 2023). Complex retroviruses such as HIV-1 are important global public health concerns, and the addition of circRNAs as new players in the host virus–arms race demonstrates the complexity of host–virus interactions(Choudhary et al., 2021). In particular, the accessory proteins encoded by HIV-1 play crucial roles in natural infection(Malim & Emerman, 2008), and how accessory proteins can also function by hijacking the host circRNA network is crucial for obtaining mechanistic insights into viral pathogenesis, which goes beyond protein‒protein interactions. The viral protein Vpr was shown to induce the expression of the *SMARCA5*-encoded circRNA ciTRAN(Bhardwaj et al., 2023). Vpr is packaged within virions and is classified as an immediate early protein that plays various roles in DNA damage, replication stalling, homologous recombination (HR) repair, and cell cycle arrest (Guenzel et al., 2014; He et al., 1995; D. Li et al., 2020; Tachiwana et al., 2006). The induction of DNA damage has emerged as one of the prominent functions of Vpr protein. For example, HIV-1 Vpr has been shown to modulate the DNA damage response (DDR) by inducing DNA damage and modulating the functions of DDR signaling proteins(Iijima et al., 2018; Schröfelbauer et al., 2007). The virological attributes suggest that Vpr is an immediate-early protein which gets incorporated within the particles during virion biogenesis(Cohen, Dehni, et al., 1990; Cohen, Terwilliger, et al., 1990; Hrimech et al., 1999). Moreover, virally delivered Vpr protein may regulate both host and viral transcription (Forget et al., 1998; Goh et al., 1998; Harman et al., 2015; Subbramanian et al., 1998). However, the connection between various activities of Vpr and its impact on cellular signaling and viral transcriptional promotion is incompletely understood.

Among various cellular signaling pathways, the transforming growth factor-beta (TGF-β) signaling is essential for regulating key cellular processes, including proliferation, differentiation, and apoptosis (Moustakas et al., 2002). This pathway is activated when TGF-β ligands bind to type I and type II serine/threonine kinase receptors, causing the phosphorylation of SMAD proteins (Nakao et al., 1997). These phosphorylated SMAD proteins then migrate to the nucleus to influence gene expression. TGF-β signaling is closely linked with DDR pathways, crucial for maintaining genomic stability. Upon DNA damage, TGF-β can modulate the DDR by affecting the expression of genes involved in cell cycle arrest, DNA repair, and apoptosis (Y. Li et al., 2019). For example, TGF-β promotes the repair of DNA double-strand breaks via the nonhomologous end joining (NHEJ) and homologous recombination (HR) pathways (Barcellos-Hoff & Cucinotta, 2014). Additionally, TGF-β signaling can induce cell cycle arrest, providing cells with more time to repair DNA damage before division. Dysregulation of TGF-β signaling is often associated with cancer, as it can influence DNA repair mechanisms and increase genomic instability, contributing to tumor progression and chemotherapy resistance (Liu et al., 2021; M. Zhang et al., 2021).

In the context of HIV-1 infection, TGF-β signaling is also known to promote virus replication in CD4+ cells and to drive the transitional effector phenotype in T cells, which combat the transcriptionally silent simian immunodeficiency virus and reduce viral reservoirs *in vivo* (Kim et al., 2024; Yim et al., 2023). Higher plasma levels of TGF-β are also connected to disease progression, impaired immune functions, increased viral replication, and CD4+ T-cell depletion, implicating TGF-β as a crucial pathogenic mediator (Dickinson et al., 2020; Kekow et al., 1990, 1991; LAZDINS et al., 1992; Wiercińska-Drapalo et al., 2004; Y. Zhou et al., 2023). However, the functional outcome of increased TGF-β signaling in cells targeted by HIV remains unclear.

Given the parallel effects of Vpr and TGF-β signaling on HIV replication, we surmised that ciTRAN induction by HIV-1 Vpr is intricately connected to TGF-β signaling.

## Results

### ciTRAN upregulation correlates with the extent of DNA damage in T cells

To obtain insights into how the expression of proviral circular RNA ciTRAN is induced by HIV-1 Vpr during infection, we employed a panel of mutants that genetically separated the activities of the accessory protein (Fig. 1A, Supplementary Fig. 1A). JTAg T cells challenged with lentiviral vector (LV) particles encapsidating Vpr protein were harvested 30 h post transduction, and the ability of indicated Vprs to induce RNaseR-resistant ciTRAN expression was assessed from total RNA using a set of divergent primers as reported earlier (Bhardwaj et al., 2023)(Supplementary Fig. 1B). The hemagglutinin (HA) tagged Vprs were detected from the cell lysates by western blotting (Fig. 1B: bottom panel). Whereas cells expressing the Vpr Q65R mutant showed comparable ciTRAN expression to that of cells without Vpr, the other mutants increased the level of ciTRAN to varying extents (4-to-6-fold) (Fig.1B). The Q65R mutant was reported to be defective in inducing the DNA damage response, in addition to arresting the cell cycle at the G2 phase, stalling the replication fork, and repressing HR(He et al., 1995; Iijima et al., 2018; Zimmerman et al., 2006). The W54R mutant was defective in binding to UNG2 but retained all the above functions, including cell cycle arrest(Guenzel et al., 2012). The S79A mutant, proficient in inducing the DDR but failed to arrest the cell cycle or stall the replication fork and HR repression (Belzile et al., 2007; Le Rouzic et al., 2007; D. Li et al., 2020), also increased ciTRAN levels. Furthermore, analysis of increase in DNA damage by tail profile via an alkaline comet assay revealed that in addition to WT Vpr, W54R, S79A, and R80A were proficient in inducing DNA damage; however, the Q65R mutant consistently failed to induce DNA damage under these conditions (Fig.1C). We also confirmed nuclear DNA damage marker γH2AX by immunofluorescence assay (Supplementary Fig. 1C) as well as its phosphorylation by western blotting (Supplementary Fig. 1D) for WT and mutant Vprs suggesting the activation of DDR pathway. Subsequent correlation analysis of endpoints from these two orthogonal assays revealed that the DNA damage function of Vpr was correlated with ciTRAN upregulation (Fig.1D). Thus, we next examined whether any DNA damaging agent can upregulate ciTRAN. Interestingly, ciTRAN expression induction by DNA-damaging agents (Etoposide (ETO) and Doxorubicin (Dox)), which cleave DNA randomly, was reminiscent of Vpr action on ciTRAN upregulation (Fig.1E). These experiments suggested that DDR may be responsible for ciTRAN upregulation, as DNA damage by Vpr, ETO or Dox can trigger ciTRAN expression.

**Figure 1.**
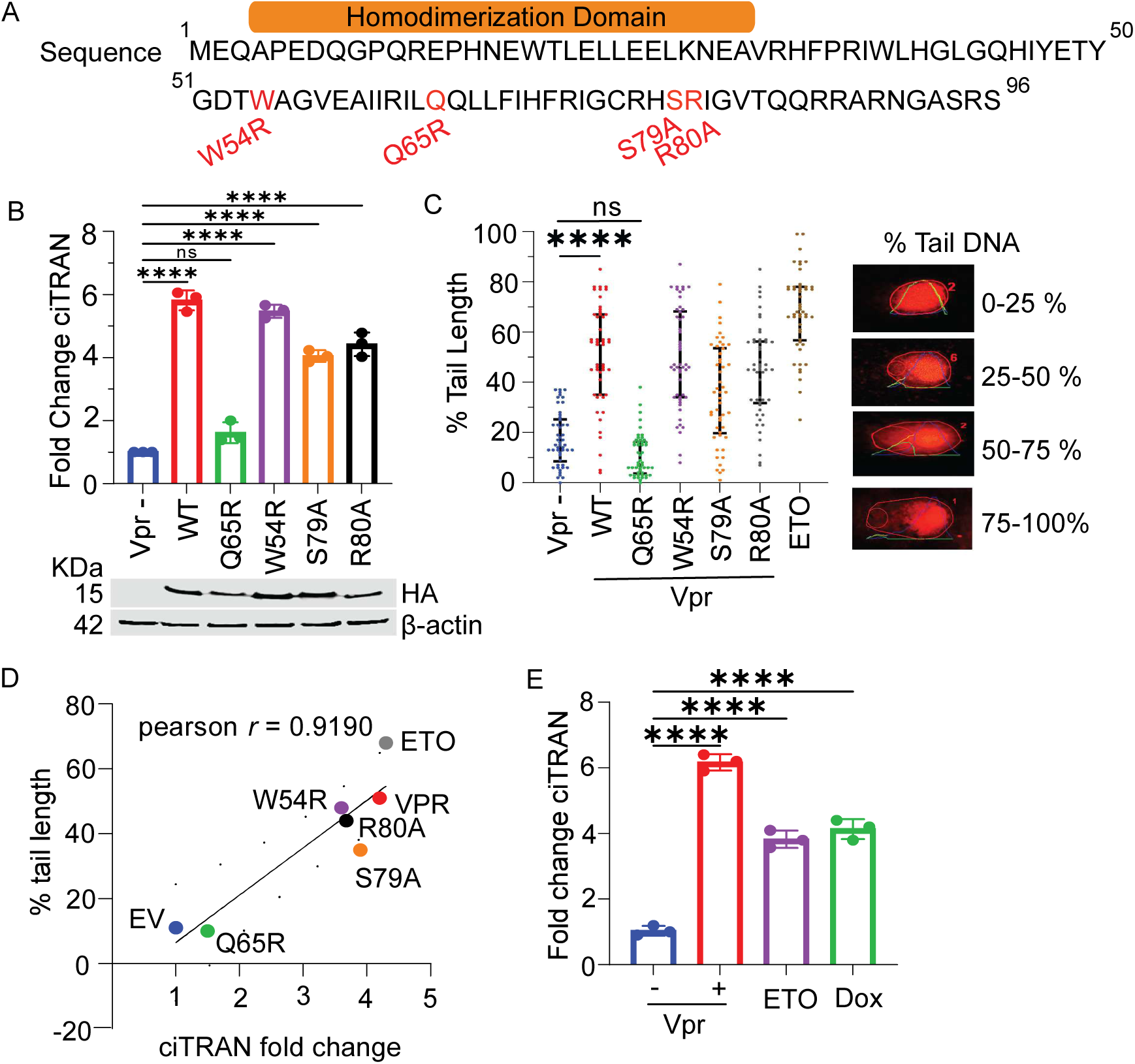
ciTRAN upregulation correlates with the extent of DNA damage caused by HIV-1 Vpr. **A**. Schematics representation of HIV-1 Vpr (uniport ID P05928), the highlighted sequences show mutation prevalence and highlighted mutants that were studied. **B**. ciTRAN levels from JTAg cells by qRT-PCR in WT or mutant Vpr expressing conditions. Vpr (-) expressing condition was used as control. Immunoblotting from transduced JTAg showing corresponding Hemagglutinin (HA)-tagged Vprs. **C**. Alkaline comet assay indicating extent of DNA damage in JTAg cells challenged with virion-packaged WT or mutant Vprs, n =50±SD. The bars represent the median with the interquartile range. Right panel, visual representation of the four degrees of damage measured by percent tail DNA. Intensity profiles, lines, and numbers on the images were automatically generated by the *OpenComet* plug-in for the software ImageJ. **D**. Correlation of percent tail-length and fold change in ciTRAN induction has been shown by plotting the mean values for each condition. **E**. ciTRAN levels after transducing JTAg cell with VLPs without or with Vpr or 6hr treatment with 10µM ETO or Dox.

### ciTRAN upregulation is independent of the cell cycle arrest and is not modulated by central players of the DDR pathway

The cell cycle arrest is a well-known function of Vpr and is dependent on the presence of its C-terminal tail. Accordingly, truncation of the C-terminal tail or substitution of the 80^th^ arginine residue with alanine prevents the ability of Vpr to arrest cells in the G2 phase (Maudet et al., 2011). Our experiments with S79A and R80A mutants suggested that cell cycle arrest may not be responsible for the upregulation of ciTRAN expression (Fig.1B). Furthermore, the W54R mutant was proficient in inducing cell cycle arrest in G2, as well as promoting DNA damage to upregulate ciTRAN expression (Fig. 1B, 1C and 2A). We thus sought to further resolve this relationship by adding specific cell cycle inhibitors that arrested JTAg T-cells in particular cell cycle stages (G2/Mi: Apigenin and G1i: CPI203).

The ciTRAN levels of the JTAg T cells challenged with the indicated inhibitors, LV-delivered WT Vpr or ETO, were then checked. While the compounds effectively inhibited the cell cycle in the respective phases (Supplementary Fig. 2A), they failed to induce ciTRAN expression (Fig.2B), additionally confirming that cell cycle arrest may not result in upregulation of ciTRAN.

**Figure 2.**
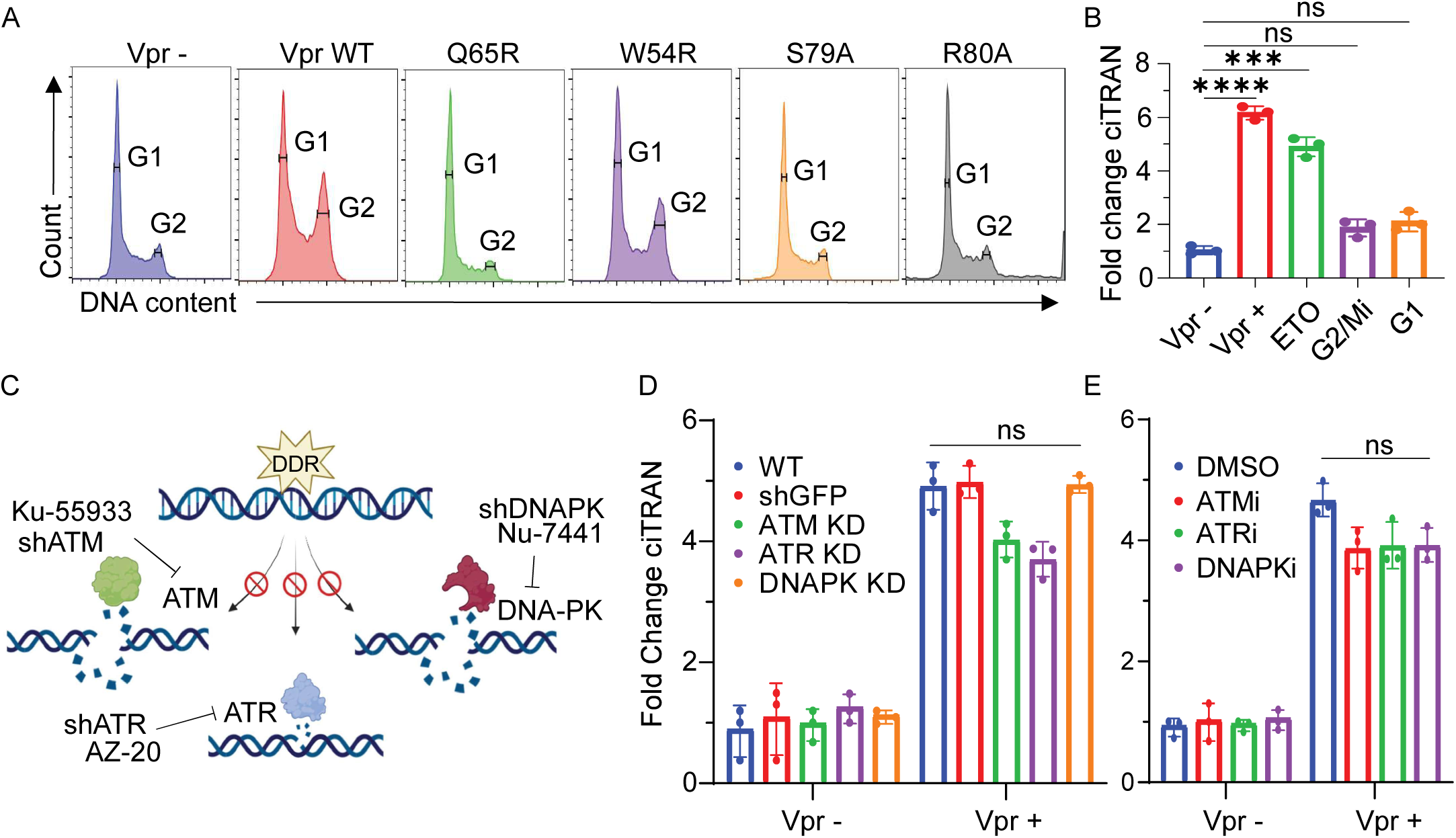
ciTRAN induction is independent of cell cycle arrest and is not regulated by central players of DDR. **A**. Cell cycle profiles from JTAg cells challenged with indicated virion-packaged Vprs. **B**. ciTRAN levels from JTAg cells challenged with Vpr+/- LVs or for 6hrs with 10µM ETO or Apigenin or CPI203. Cells were harvested after 24 hours for quantification of ciTRAN by qRTPCR, GAPDH served as control for normalization. Central players of DDR viz ATM/ ATR/DNAPK (**C**) were inhibited either by lentiviral RNAi (**D**) or specific small molecule inhibitors (**E**) for 6 hours to assess the impact of those on ciTRAN levels in Vpr +/- conditions. Vpr containing LVs were used to deliver the protein to JTAg wild type (WT) cells or shGFP or shATR or shATM or shDNAPK stably expressing cells. After 24 hours, the intracellular levels of ciTRAN were assessed by qRT PCR.

Next, because DNA damage appeared central to the induction of ciTRAN expression, we asked whether the central players of the DDR pathway, such as ATM, ATR and DNAPK, which are activated upon DNA damage, are responsible for ciTRAN upregulation (Fig.2C). Accordingly, JTAg T cells were selected by puromycin after LV transduction for shRNA-mediated knockdown (KD) of ATM, ATR, and DNAPK. A nonrelevant shRNA targeting the GFP cell pool was also generated parallelly. These shRNA-expressing cells were subsequently challenged with LV particles encapsulating Vpr. While specific shRNA expression consequently reduced the mRNA levels of ATM, ATR and DNAPK (Supplementary Fig. 2B), they had an insignificant effect on the levels of ciTRAN induced by Vpr (Fig.2D). This was independently confirmed by using select inhibitors for each of the central DDR players: Ku-55933 (ATM), AZ-20 (ATR), and Nu-7441 (DNAPK) (Fig.2E). Taken together, these results suggest that cell cycle arrest, the most prominent feature of Vpr, is dispensable for ciTRAN upregulation. In addition, the experiments further indicated that the central players of the DDR pathway may not regulate ciTRAN expression.

### ciTRAN upregulation by Vpr is orchestrated via the TGF-SMAD axis

Considering that DNA damage is central to the transcriptional induction of ciTRAN, we hypothesized that the downstream pathways activated in response to DNA damage by Vpr or any DNA damaging agent are involved in ciTRAN upregulation. This is because DNA-damaging agents such as etoposide and doxorubicin could induce ciTRAN expression in the absence of infection (Fig. 1E). Notably, the upregulation of ciTRAN as well as its parental mRNA (SMARCA5) by WT Vpr as well as other mutants, excluding Q65R, was consistent (Supplementary Fig.3A), suggesting a promoter level induction of the transcript. Emerging evidence indicates that TGF-β signaling is activated in response to treatment with DNA-damaging agents, such as doxorubicin and etoposide, in cancer patients as well as in cell lines and can affect the clinical outcomes of patients undergoing chemotherapy (Y. Li et al., 2019; Liu et al., 2021). Moreover, our recent study reported that TGF-β signaling can also get activated downstream of sequence-specific DNA cleavage via programmable nucleases such as Cas9 (Mishra et al., 2022). We thus hypothesized that the TGF beta pathway downstream of Vpr expression is an intermediate signaling module potentially required for the upregulation of ciTRAN.

To test this, we performed reporter assays in HEK293T cells using the TGF-β -sensitive luciferase plasmid (SBE-Luc; Fig.3A top panel) to determine whether Vpr can induce TGF-β signaling. In this construct, luciferase expression is driven by a promoter carrying SMAD binding element (SBE). Along with SBE-Luc, the HEK293T cells were transfected with either empty (EV) or DNA damage-proficient Vpr (WT) or DNA damage-deficient (Q65R) vectors. In addition, we also transfected cells in separate wells with SBE-Luc along with a *gCXCR4*-Cas9-encoding plasmid as control, which generated sequence-specific DNA breaks guided by gRNA specific to *CXCR4* (Mishra et al., 2022). Recombinant TGF-β served as positive control in this assay. Notably, the luciferase reporter assay indicated that WT Vpr expression, in addition to *gCXCR4*-Cas9 expression, is associated with the activation of TGF-β signaling (Fig. 3A). However, the DNA damage-deficient Vpr (Q65R) failed to induce luciferase expression, emphasizing the specificity of the assay and, for the first time, the ability of WT Vpr to mediate the activation of TGF signaling (Fig.3A). Strikingly, we found that *gCXCR4*-Cas9-induced sequence-specific DNA cleavage (Fig. 3B) or the addition of recombinant TGF-β (Fig. 3C) in HEK 293T cells was also sufficient to upregulate ciTRAN levels almost equivalent to those of WT Vpr expressing cells. Additionally, in JTAg T cells, the TGF-β signaling effectors SMAD2/3 were also examined for activating phosphorylation, and in agreement, we found that cells transduced with LV particles containing WT Vpr presented elevated levels of intracellular p-SMAD2/3, which expectedly, were reversed by addition of TGFBR1 inhibitor Repsox (Fig.3D).

**Figure 3.**
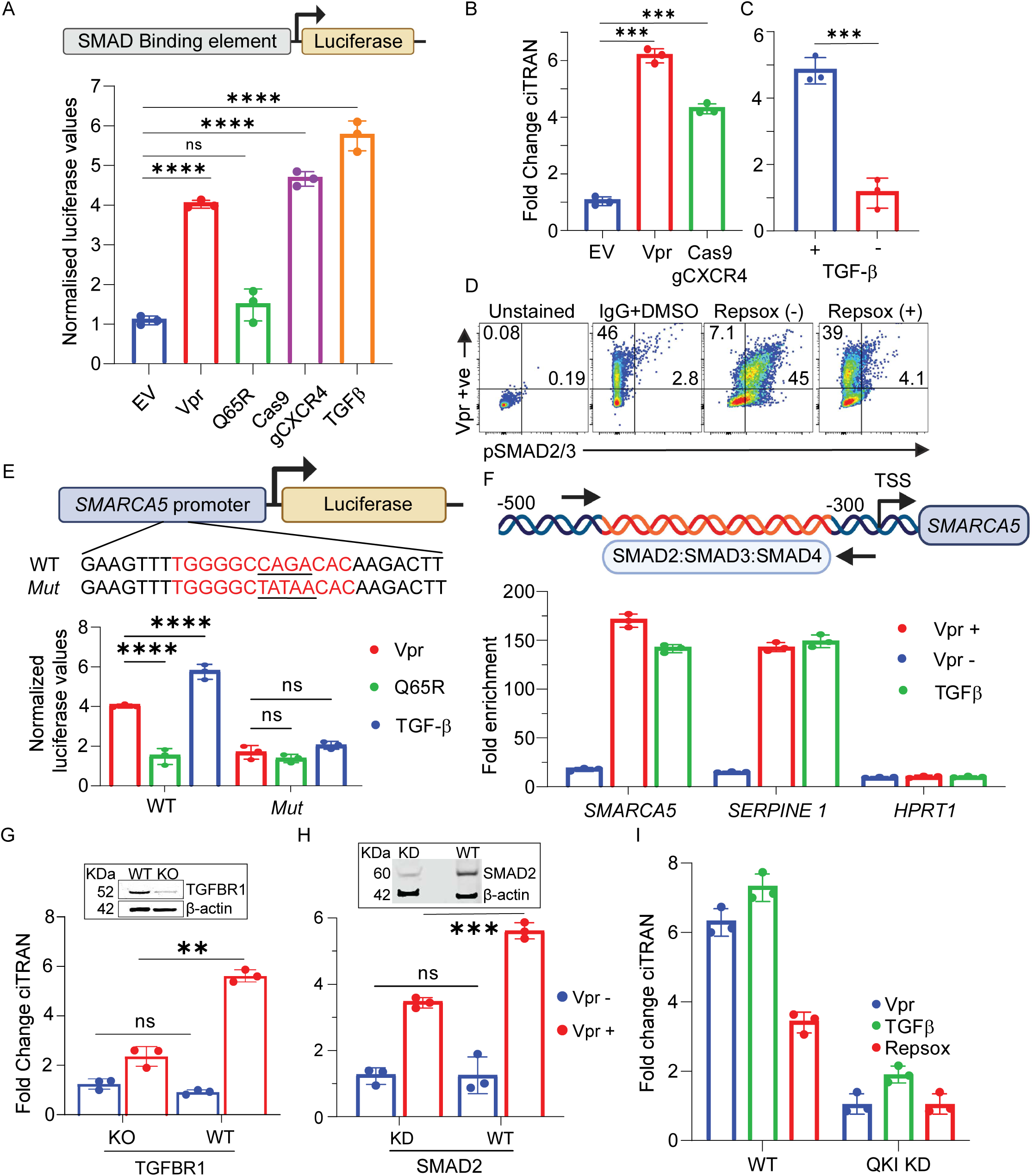
Vpr promotes TGF signaling which subsequently orchestrates ciTRAN induction. **A**. HEK293T cells transfected with SMAD-sensitive luciferase reporter (SBE Luc) along with either empty vector (EV), or plasmids expressing Vpr or cas9 and luciferase assay was performed to measure extent of SMAD-dependent luciferase expression. Recombinant TGF β (5ng/ml) was used as control for activating TGF β-SMAD signling and concomitant luciferase expression from SBE Luc construct. Data were normalised by total protein amount (SD+/-). ciTRAN levels (as in Fig2D) from: (**B**) HEK293T cells transfected with empty vector (EV) or Vpr or Cas9 expressors, and (D) JTAg cells challenged with TGF-β(10ng/ml), at 24 hours. GAPDH serves as control (n=3 ±SD). **C**. Flow cytometry analysis of JTAG cells transduced with GFP lentivirus encapsidated with Vpr protein. The unstained and untransduced JTAG cells were used as control. The population expressing pSMAD2/3 was detected by staining with Alexa 633 labeled secondary Ab (APC channel) added following the incubation of target cells with either isotype control (IgG) or p-smad2/3 antibody. The GFP was used as indirect marker for Vpr expression. The specificity of smad phosphorylation in these assays was further confirmed by a TGFR1/Alk5 inhibitor Repsox. **E**. Schematics of putative WT SMAD2/3/4 or Mut-SMARCA promoter regions (detailed in methods) inserted upstream to the Luciferase gene in pGL3-basic. HEK293T cells transfected with WT or Mut SMARCA5 promoter plasmids along with Vpr expressing or Vpr Q65R expressing or empty vector. Luciferase readout was taken after 24 hours. Recombinant TGF-β served as positive control ( n=3 ±SD). **F**. Top panel: schematics of putative SMAD binding motifs in SMARCA5 promoter at the indicated locations. The arrows show the position of oligos (used in ChIP assay) binding to the promoter region with respective positions indicated upstream to the transcription start site. Bottom panel: Enrichment of SMARCA5 promoter fragment by SMAD2/3 specific antibody examined by ChIP-qPCR assay. IgG served as antibody specificity control. (n=3 ±SD) SPERPINE1 was used as positive control whereas HPRT1 intronic region was taken as negative control for SMAD2/3 recruitment. **G**. ciTRAN levels assessed after 30h challenge of Vpr+/- lentiviral particles in JTAG cells by qRTPCR from: (**G**) Wild-type (WT) or TGFRI knockout (KO) cells and (**H**) control shRNA (SMAD2 WT) or SMAD2-specific (SMAD2 KD) shRNA expressing cells, selected by puromycin (n=3 ±SD). Data was normalised to actin mRNA levels. Corresponding immunoblots of TGF-β RI and SMAD2 from G and H in the insets with respective loading shown by probing β-actin from the same cell lysates. **I.** Effect of QKI knockdown on the upregulation of ciTRAN by Vpr, TGFβ and the sensitivity to Repsox.

Intriguingly, analysis of the promoter sequence of *SMARCA5,* the parental gene that encodes ciTRAN, revealed the presence of a SMAD binding consensus motif, which suggested the plausible recruitment of crucial TFG-β signaling effectors (SMAD2/3) to promote circRNA expression (Supplementary Fig.3B). To validate this in silico observation experimentally, we selected a promoter fragment (1157 bp) of SMARCA5 and cloned it upstream of the luciferase gene. Notably, we found that SMARCA5 promoter-driven luciferase expression was dependent on either the expression of Vpr or the challenge of cells with recombinant TGF-β (Fig.3E). Furthermore, the DNA damage-defective Vpr mutant or the promoter variant (Mut-*SMARCA5*) that lacked the predicted SMAD binding element could not induce luciferase reporter expression under these conditions (Fig.3E), suggesting the dependency on the DNA damage and subsequent TGF-β canonical signaling activation for reporter expression. To further independently confirm the SMARCA5 promoter occupancy by SMAD2/3 and to rule out indirect effects on luciferase expression from the transfected plasmid, we examined the endogenous *SMARAC5* promoter (Fig. 3F, top panel) in chromatin immunoprecipitation (ChIP) assays using SMAD2/3 antibody. Compared with the Vpr minus condition, the antibody specific to SMAD2/3 enriched the SMARCA5 promoter sequence 169-fold from the lysates of cells challenged with Vpr-loaded LV particles. Recombinant TGF-β was used as a positive inducer of TGF-β signaling. This altogether indicates, for the first time, that Vpr-induced TGF beta signaling occurs in physiological settings and that SMARCA5/ciTRAN expression is regulated by SMAD2/3, classical mediators of TGF-β signaling (Fig.3F, bottom panel).

Additionally, the specificity of the assay was also confirmed by selecting a known downstream effector gene, SERPINE1, as a positive control and HPRT (intron) as a negative control (Fig.3F, bottom panel). In conditions where TGFβsignaling was promoted, either by DNA damage or by direct engagement of TGF receptors by the recombinant ligand, we found that the chromatin region of SERPINE but not that of HPRT was enriched, indicating the specificity of the assays performed.

Next, we investigated major regulators of TGFβ signaling, starting with CRISPR targeting of TGFBRI to determine whether the major receptor is required for ciTRAN upregulation by Vpr. Indeed, genetic ablation of the TGFBR1 locus affected the ability of Vpr to upregulate ciTRAN; in the gene-edited bulk population, the expression difference was reduced to a mere 1.5-fold from 6-fold (Fig.3G). Western blotting correspondingly confirmed the loss of the TGFRI signal in the gene-edited bulk population (Fig.3G, inset). The degree of TGF dependency was also evaluated by knocking down the primary TGF effector SMAD2 (Fig.3H and inset) and pharmacologically targeting TGFBR1 via Repsox (Supplementary Fig.3C). As expected, we observed a reduction in the magnitude of ciTRAN induction by Vpr in SMAD2-knockdown cells as well as in Repsox-treated cells. However, notably, regardless of TGF-β signaling activation by Vpr or recombinant TGF-β, ciTRAN upregulation still required the QKI protein (Fig.3I and Supplementary Fig.3D). Taken together, these experiments provide new insights into the positive regulation of TGF-β signaling by HIV-1 Vpr and the upregulation of circRNA ciTRAN expression by Vpr-induced TGF-β signaling.

### HIV replication promoted by ciTRAN is decreased by TGF-β signaling inhibition

We next sought detailed insights into the regulation of ciTRAN expression by TGF-β signaling and to determine the contribution of upregulated ciTRAN to the formation of functional RNA polymerase-II (RNAPII) transcription complex on HIV-1 LTR promoter. Accordingly, we examined SRSF1, RNAPII, and Tat occupancy via ChIP assays in conditions where Repsox or DMSO inhibited TGF-β signaling. A strong interaction of SRSF1 with the HIV-1 TAR locus was observed in conditions where the cells were challenged with Repsox according to the ChIP assay (Fig. 4A), which is consistent with our previous study (Bhardwaj et al., 2023) that a competing scenario between ciTRAN and SRSF modulating HIV-1 proviral transcription (Fig. 4A). These findings were also consistent with the reduced association of RNAPII and Tat with the TAR region in repsox treated cells (Fig. 4A). TGF-β signaling inhibited by Repsox concurrently inhibited ciTRAN expression (Supplementary Fig. 3C), which negatively impacted the steady-state levels of provirus-encoded HIV RNA as examined from nuclear and the cytoplasmic compartments (Fig. 4B and 4C). Correspondingly, decreased replication of HIV-1 in the presence of Repsox in E6.1 T cells was also observed (Fig. 4D). Overall, the levels of viral transcripts and gene expression were impacted by Repsox treatment, as reflected by the percentage of reporter (zsGreen) positive cells detected by flow cytometry and the magnitude of zsGreen expression (Fig. 4D). Moreover, a CRISPR assay with the TGFBRI primary receptor in JTAg T cells and subsequent infection by zsGreen-expressing HIV-1 was performed to confirm this phenomenon independently (Fig-4E).

**Figure 4.**
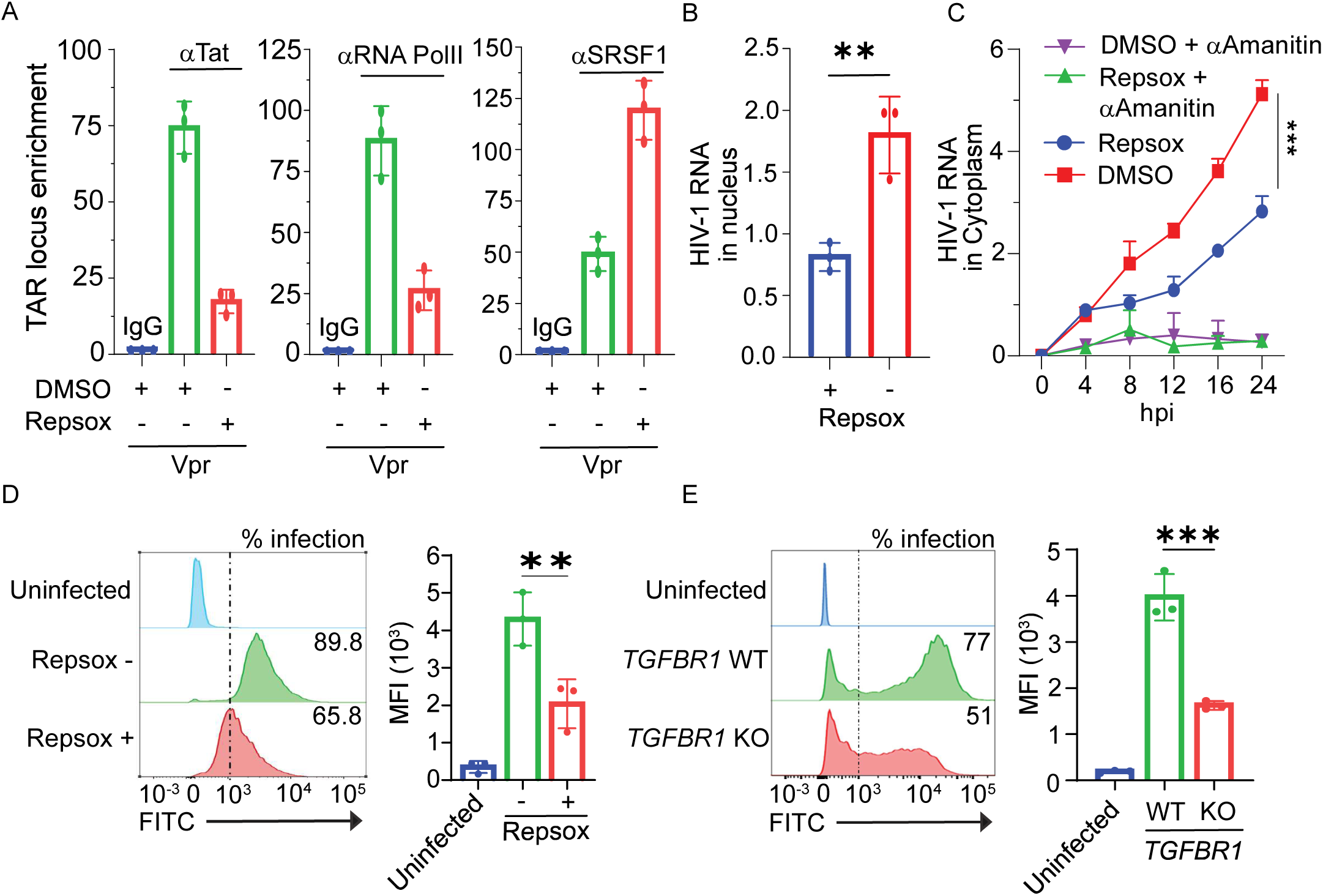
TGF-β signalling inhibition restricts HIV replication by downregulating ciTRAN. **A**. TAR locus enrichment and analysis by ChIP-qPCR assay. JTAG cells were challenged with lentiviral particles encapsidating Vpr and were treated with DMSO or Repsox (10µM). The ChIP was performed after 24 hours using RNA polII-, SRSF1-, and TAT-specific antibody. The data was normalised to TAR region enriched in IgG precipitated fractions (n=3 ±SD). Transcription from HIV-1 provirus analysed by gag-specific qRT-PCR in Repsox treated and untreated conditions: (**B**) Nuclear run-on assay to measure the levels from infected cell nucleus and (**C**) cytoplasmic fraction collected across indicated time points to measure gag RNA levels within the cytoplasm (α-Amanitin served as a control). The data was normalised in each condition to GAPDH mRNA. **D, E.** The frequency (expressed as percent positive cells) and the magnitude of HIV-1 gene expression (expressed as Mean Fluorescence Intensity of HIV-1 encoded zsGreeen reporter (MFI)) captured by flow-cytometry after 24 hours of infection of E6.1 treated +/- repsox (**D**), and JTAg cells lacking functional TGFBR1 (**E**) n = 3; ±SD.

### TGF-β signaling inhibition suppresses transcription from transmitted founder virus LTRs

We next investigated whether small-molecule inhibitors of TGF-β signaling can reduce the transcription from LTRs of transmitted founder (TF) viruses by depleting ciTRAN levels. Accordingly, two distinct small molecules (Repsox and SB525334) known to target TGF-β signaling were used to challenge HEK293T cells transfected with molecular clones of the indicated TF viruses and ciTRAN levels were captured by qRT PCR. We observed decreased ciTRAN levels in these cells (Fig.5A), and correspondingly reduced transcription from LTRs of the TF viruses (Fig. 5B, SRSF ectopic expression was used as control in these reporter assays as reported earlier(Bhardwaj et al., 2023)), suggesting the conserved nature of these signaling cascades in the replication of clinically relevant isolates. Furthermore, we investigated whether monocytic lineages (such as THP-1), where Vpr profoundly impacts viral gene expression(Subbramanian et al., 1998), are also sensitive to Repsox treatment. Indeed, Repsox treatment decreased the expression and frequency of zsGreen-positive THP-1 cells (zsGreen encoded by the provirus in THP-1 cells) (Fig. 5C).

**Figure 5.**
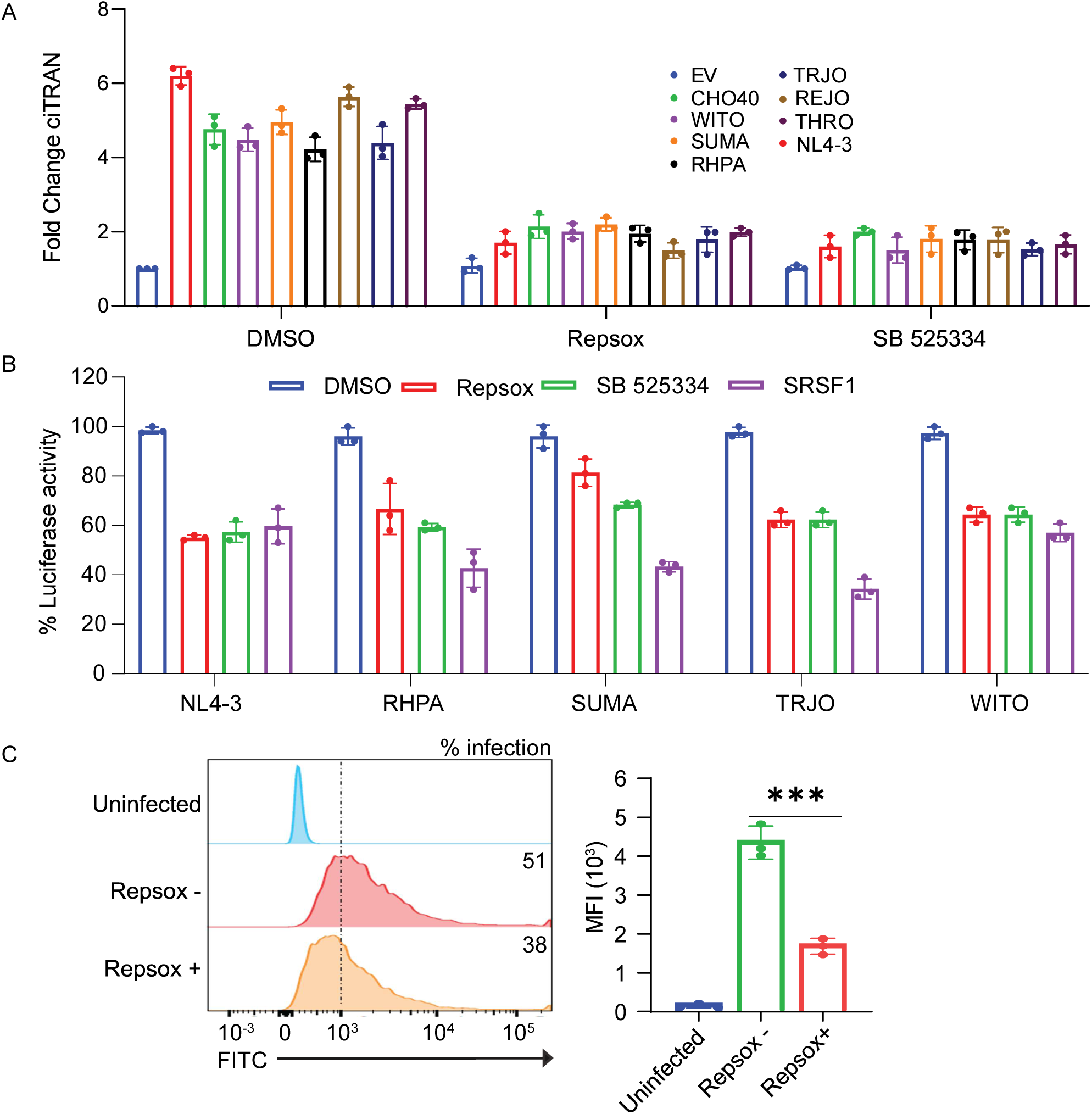
Effect of TGF signaling inhibition on TF virus transcription and HIV replication in monocytes. **A.** ciTRAN levels from TF virus molecular clones transfected HEK293T cells challenged with indicated inhibitors. **B**. Transcription from LTRs of different TF viruses and its sensitivity to inhibitors. SRSF1 ectopic expression was used as control. **C**. HIV-zs green replication in THP-1 monocytes in indicated conditions assessed by Flow-cytometry.

### The role of TGF-β signaling in natural infection and the inhibition of HIV replication by Repsox in primary cells

Our analysis of the publicly available RNA seq data (from Ref# (Y. Zhou et al., 2023)) revealed a significant upregulation of TGFb1 in HIV-1 infected patients as compared to healthy counterparts (n=6, Fig. 6A). Intriguingly, analysis of an independent additional dataset (PRJNA824711) of HIV patients including regular progressors (Regular), long-term nonprogressors (LTNP), and non-controllers (LNC) analyzed for TGF superfamily cytokines (TGF-β1-3; Supplementary Fig.4B) revealed a significant TGF-β1 transcript expression in regular but not in LNCs or LTNPs (n=4, Supplementary Fig.4A). The significantly increased TGF-β1 levels in these patients (Supplementary Fig.4A) positively correlated (r=0.8412) with the plasma viral load (Supplementary Fig.4C) suggesting towards significance of the increased TGF-β1 in the disease progression and maintain higher plasma viral load. Thus, we next sought to understand the importance of TGF-β signaling for HIV replication in the context of ciTRAN using primary CD4+ T cells enriched to purity by positive selection (Supplementary Fig.4D) from pooled PBMCs from three donors. We checked the TGF-β1 isoform expression in primary CD4+ T cells; indeed, TGFB1 was upregulated in Vpr-treated samples (Fig. 6B). Moreover, the extent of TGF-β1 signaling concorded with ciTRAN expression in primary cells treated with LVs carrying Vpr (Fig. 6C), and ciTRAN expression was sensitive to repsox treatment regardless of Vpr expression (Fig. 6C). Notably, reduced ciTRAN levels upon challenge by repsox were not associated with reduced viability of primary CD4+ cells in these conditions (Supplementary Fig.4E).

**Figure 6.**
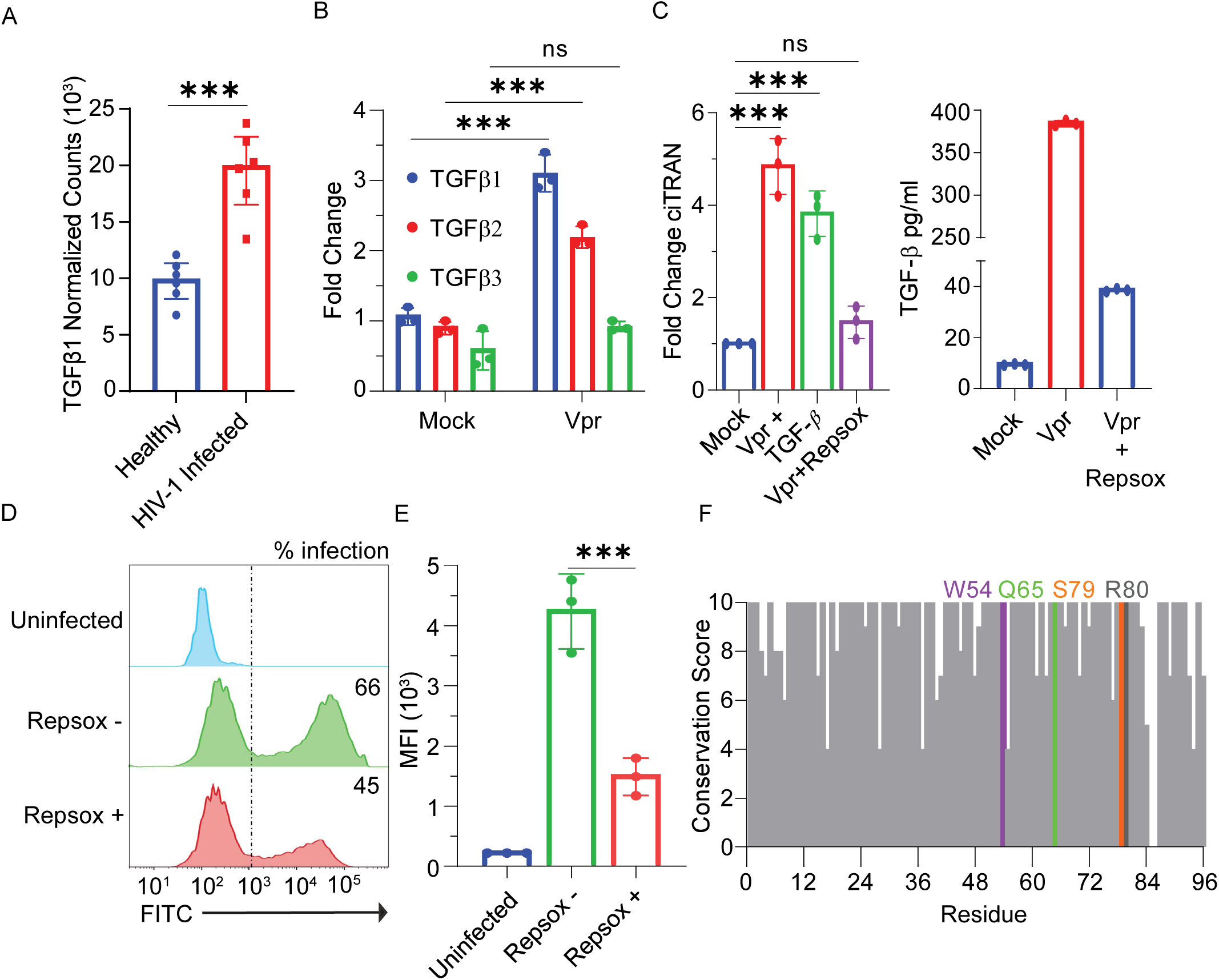
TGF-β signaling in natural infection and inhibition of HIV replication by repsox in primary human CD4+ cells. **A**. Analysis of TGF-β transcripts from PBMCs of healthy and HIV-positive individuals. **B**. Analysis of TGF-β superfamily members from human primary CD4+ cells by qRT-PCR in Vpr +/- conditions. **C**. Primary CD4+ cells were infected with HIV-1 NLBN zsgreen repsox (+/-), conditioned media was used as mock and recombinant TGF-β was used as positive control. Subsequently, cells were harvested for the ciTRAN analysis with q-RTPCR n = 3; ±SD. **D**. ELISA for total TGF-β was performed from supernatant of primary CD4+ cells infected with HIV-1 zsGreen (+/-) Repsox for 24 hours in a serum-free medium. **E**. Effect of Repsox on HIV-1 zsgreen replication in primary CD4+ cells. MFI, mean fluorescence intensity, n = 3; ±SD. **F.** Conservation of indicated amino acid residues in Vpr sequences analyzed from the LANL database.

Furthermore, an ELISA specific for detecting cell-free TGF-β confirmed that Vpr strikingly induced TGF-β expression (380pg/ml) and that was reversed by addition of repsox to 39 pg/ml (Fig-6D). Specifically, in this case, the advanced RPMI was used to grow the cells containing minimal FBS (1%) to exclude the possibility of TGF-β1 being detected from a bovine source in ELISA. Finally, we checked single-cycle HIV replication in primary cells challenged with Repsox or DMSO. In agreement, the impact on the proportion of positive cells, as well as the magnitude of reporter expression, was negatively influenced by Repsox treatment (Fig. 6E). The proportion of cells positive for infection was reduced from 66% to 45%, and the reporter expression was reduced from 4.2-fold to 1.4-fold after the cells were treated with repsox, confirming the inhibition of HIV-1 gene expression by the TGF-β small molecule inhibitor in primary cells.

## Discussion

Here, we demonstrate an intriguing connection between HIV-1 Vpr-mediated DNA damage and the consequent activation of TGFβ signaling leading to upregulation of a proviral circRNA ciTRAN. Vpr is a unique protein encoded by primate lentiviruses known to promote viral replication (Forget et al., 1998; Malim & Emerman, 2008), and the selective pressure exerted to maintain the functional copy of a gene implicates its relevance for natural infection (Ali et al., 2018; Goh et al., 1998). Notably, the glutamine residue at the sixty-fifth position is remarkably conserved across HIV-1 Vpr sequences we analyzed (Fig.6F). This implies the selective pressure exerted to achieve DNA damage and plausibly TGF-β signaling activation functions *in vivo*. Vpr functions are now envisaged beyond regulating protein-protein interactions, as exemplified by the co-option of a circular RNA. SRSF1, a constitutively expressed protein known to impact viral gene expression significantly (Paz et al., 2014), was recently shown to be antagonized by Vpr by promoting the expression of ciTRAN(Bhardwaj et al., 2022). However, the molecular mechanism of ciTRAN induction remained elusive. Clearly, the induction of ciTRAN by HIV-1 Vpr is actively achieved by the cascade of signaling events mediated by the accessory protein, as demonstrated in this study. The experiments from various cell lines and primary CD4^+^ cells reveal that Vpr induces DNA damage that precedes the activation of TGFβ signaling, which subsequently promotes infection by inducing ciTRAN expression. Furthermore, we showed that TGFβ inhibition by orthogonal means significantly reduced ciTRAN expression during infection, suggesting an effect on viral replication independent of proteins involved in the DDR and cell cycle arrest. This was relevant to check because HIV-1 was known to exploit the Fanconi anemia pathway for its replication and Vpr is known to arrest cycling cells(Fu et al., 2022; Goh et al., 1998).

While TGF-β signaling has been implicated in HIV replication (LOTZ & SETH, 1993; Yim et al., 2023), admittedly, the direct link between TGF beta induction, viral protein involvement, and circRNA levels was not evidenced from the RNA-seq data from patients analyzed in this study. This may be because the datasets were generated from the peripheral blood cells that lacked information on circRNAs and expression of specific viral genes associated with the infected target cell population (Y. Zhou et al., 2023). circRNAs can be further explored via improved sequencing protocols that allow analysis of such non-coding RNAs from virally infected cell populations. In this study, experiments, including those using primary CD4^+^ T cells and orthogonal perturbations, were conducted to explore this link and show for the first time that TGFβ signaling is induced by the viral accessory protein Vpr which can positively influence HIV-1 gene expression via ciTRAN upregulation. Moreover, this study shows how circRNA functions can be targeted by small molecules that directly impact circRNA levels to suppress viral replication. Accordingly, the use of repsox as an adjunct to current antiretrovirals during acute infection may not only help circumvent the immunosuppressive activities of TGF-β1 but also reduce the viral burden by inhibiting circRNA expression (Gopalakrishnan et al., 2021; O’Reilly et al., 2014) and prevent the formation of reservoirs in nonimmune cells(Chinnapaiyan et al., 2019).

Considering the DNA damaging agents routinely used in cancer treatment can also promote ciTRAN expression, our findings have implications in the clinical management of HIV-positive cancer patients and for the treatment of cancers that are independently shown to have upregulation of ciTRAN (circSMARCA5) (Xu et al., 2020). Furthermore, whether IL6 is involved in an additional positive feedback mechanism that promotes TGFβ signaling in HIV infection(Xiao et al., 2005) remains an intriguing possibility. Vpr also targets IL6 and Tet2, which are positive and negative regulators, respectively, of HIV replication, and how this interplay increases TGF-β signaling to promote HIV transcription through ciTRAN remains to be explored. The importance of targeting TGF signaling activation by HIV-1 Vpr is also stressed by the fact that TGF-β signaling can lead to pathologic concentrations of IL6 (D. Zhou et al., 1991) and the development of Kaposi sarcoma (Roth, 1993). Taken together, the findings of this study highlight the induction of TGF-β signaling by HIV-1 Vpr to promote viral replication via ciTRAN and provide a small molecule-based strategy to circumvent this phenomenon.

## Methods

### Cell culture

Jurkat E6.1 (ATCC), THP-1 (ATCC), and Jurkat TAg (JTAg)(Rosa et al., 2015)cell lines were cultured in Gibco RPMI supplemented with 10% fetal bovine serum (Certified, heat-inactivated serum from Gibco).HEK293T (ECACC), cells were maintained in 10 % FBS containing Dulbecco’s modified Eagle medium with 2mM L-glutamine. The cells were maintained in a humidified 5% CO2 incubator at 37°C

### Lentiviral Vector (LV) particle production

Lentiviral Vector particles were produced by transfecting HEK293T cells using calcium phosphate transfection reagent in a 10cm plate, pScalps ZsGreen, pMD2.G (VSVG) along with 2μg Vpr WT or other mutants expressing construct (pCDNA-HAVprs) or vector alone. After 12-15 h post-transfection, the medium was replaced with 2% FBS containing DMEM. After 48h, the virus-containing supernatant was collected at 500 x g for 5 min and filtered using 0.2mm Syringe filter.

### Virus production and infection

For infection, the virus was produced from HEK293T producer cells using the calcium phosphate method by co-transfecting 7 μg NLBN zsGreen Env defective and Nef defective (HIV-1 zsGreen reporter) and 1 μg pMD2.G encoding VSVG glycoprotein and was limited to single-cycle replication. NLBN zsGreen was a kind gift from Prof. Massimo Pizzato, which is a derivative of NL4-3 Env-Nef-described in (Pizzato et al., 2007)(Rosa et al., 2015), where env region is deleted, and zsGreen is cloned in place of Nef. The virus-containing culture supernatant was collected after 48h post-transfection, centrifuged at 300xg for 5 min, and filtered using a syringe filter of 0.22μ. The virus was quantified using an SGPERT assay(Pizzato et al., 2009), and the multiplicity of infection (MOI) was calculated by infecting HEK293T cells. Primary CD4+T cells were infected at MOI=5.

### SYBR green I-based product-enhanced reverse transcriptase (SGPERT) assay

Following the collection and filtration of the virus-containing media, 5 μL of the virus was lysed with 5 μL of lysis buffer containing 0.25% Triton X-100, 50 mM KCl, 100 mM Tris-HCl (pH 7.4), and 40% glycerol.The reaction was incubated for 10 min at room temperature and then mixed with 90 μL of 1× core buffer, which consisted of 5 mM [NH4]2SO4, 20 mM KCl, and 20 mM Tris-HCl at a pH of 8.3. Further,10 μL of the mixture was combined with 10 μL of 2× reaction buffer containing 5 mM [NH4]2SO4, 20 mM KCl, 20 mM Tris-Cl (pH 8.3), 10 mM MgCl2, 0.2 mg/mL bovine serum albumin (BSA), 1/10,000 SYBR green I, 400 μM dNTPs, 1 μM forward primer(5ʹ-TCCTGCTCAACTTCCTGTCGAG-3ʹ), 1 μM reverse primer (5ʹ-CACAGGTCAAACCTCCTAGGAATG-3), and 7 pmol/mL MS2 RNA to quantify RT units by qPCR analysis (Pizzato et al., 2009).

### RNA isolation, cDNA and q-PCR

For RNA isolation, cells were lysed using TRIzol (Invitrogen, Thermo Fisher Scientific, USA) and phase-separated using chloroform. Samples were treated using DNaseI (thermo) to remove DNA contamination. cDNA first-strand was synthesized by random hexamers or Oligo(dT) primers using Revert aid (H minus). SYBR green-I-based qPCR was performed for expression quantification, and data were represented as log2 fold change.

### Ribonuclease R treatment

For Validation of ciTRAN (circ-SMARCA5) we treated the RNA pool with RNAseR. 1 μg RNA was incubated with RNaseR (1unit) at 37°C for 30 min. RNaseR was then inactivated at 90°C for 10 minutes. RNA was then eluted using a Zymo clean and concentrator kit for RT-qPCR to quantify respective RNAs.

### Alkaline comet assay

A modified alkaline comet assay procedure was followed as previously described(Maiuri et al., 2017).JTag cells were resuspended at 4 × 10^4^ cells/mL in 1× PBS after infection with VPR mutants packaged Lentiviral particless. Cells were mixed with low melting agarose (1%) (VWR Life Science) at a 1:5 ratio and spread over the slide pre-coated with 1% normal agarose. Slides were dried at room temperature for 10 min and immersed into alkaline lysis buffer (10 mM Tris-HCl, pH 10, 2.5 M NaCl, 0.1 M EDTA, 1% Triton X-100)for 4 h and then in the alkaline running buffer (0.3 M NaOH, 1mM EDTA) for 30 min and finally electrophoresed at 300 mA for 30 min, all done at 4°C. Samples were washed thrice with double-distilled water (ddH2O) and fixed in 70% ethanol at 4°C. Slides were dried for 2 h at room temperature and stained with EtBr solution (2 μg/mL in water). Images were acquired using a Zeiss Apotome fluorescence microscope (Carl Zeiss) using a 20× objective lens, and the tail moments were quantified using the OpenComet (https://cometbio.org/). For each condition, at least 50 cells were analysed using OpenComet plugin(Gyori et al., 2014)

### Cell cycle Analysis

For cell analysis JTag cells treated with lentiviral particles carrying VprWT or mutants or treated with, G2/M inhibitor(Apegenin), G1 inhibitor(CPI 203).Cells were harvested after 48 hours and fixed in ice-cold 70% ethanol for 2 hours at 4°C. Cells were then washed in 1XPBS and stained with a staining solution containing PI(50µg/ml)and RNAseA (50µg/ml) for 30 minutes and proceeded for their DNA content analysis using flow cytometry (FACS BD AriaIII). The fluorescence of 10,000 cells was analysed using FLOWJO software.

### Smad reporter assay

To check the effect of ciTRAN induction upon inhibition of the TGFBRI pathway, we used the SMAD reporter plasmid SBE4-Luc (16495; Addgene). This plasmid contains the luciferase reporter gene under the SMAD binding elements, which are expressed under TGFBRI-mediated signaling.

HEK293Tcells were seeded in a 24-well plate 24 h before transfection. Next, cells were transfected with SBE4-Luc, , pcDNAHAvpr and empty plasmids. The transfected cells were first lysed using 100 μL lysis buffer (1% Triton X-100, 25 mM tricine [pH 7.8], 15 mM potassium phosphate, 15 mM MgSO4, 4 mM EGTA, and 1 mM DTT) for 20 min at room temperature. Luminescence readings were obtained using a Spectramaxi3X plate reader, by mixing 50 μL cell lysate supernatant with 50 μL substrate buffer (lysis buffer with 1 mM ATP, 0.2 mM D-luciferin). Data was normalized to Bradford readings of the cell lysate.

### *SMARCA5* promoter constructs and luciferase assay

To Check the TGF-β signaling affecting the SMARCA5 transcription, we generated the SMARCA5 promoter-luciferase construct; human SMARCA5 (transcript, NM_003601) promoter sequence was retrieved from the Eukaryotic Promoter Database)(Meylan et al., 2020). The SMARCA5 gene promoter from −1452 bp to −343 bp upstream of the TSS at +1 was amplified by PCR using Jurkat E.6.1 T cell genomic DNA as a template and cloned into a pGL3-Basic expression vector (Promega, E1751). SMARCA5 promoter construct was cloned between the SmaI sites. For the Mut smarca5 generation, SMAD2:SMAD3:SMAD4 binding sites were retrieved from the Jasper Motif database(Khan et al., 2018), and site-directed mutant construct was prepared using oligonucleotides with mutations in the SMAD site present (GC<u>CAGA</u>C to GC<u>TATA</u>AC). Primer details are given in Table S1. Mutations were confirmed using Sanger sequencing. For luciferase assay, HEK293T cells were seeded in a 24-well plate 24 h before transfection. Next, cells were transfected with SMARCA5 WT promotor-Luc or SMARCA5Mut promotor-Luc with pcDNAHA-VprWT or pcDNAHAQ65Rvpr or TGF-β as positive control. The transfected cells were first lysed using 100 μL lysis buffer (1% Triton X-100, 25 mM tricine [pH 7.8], 15 mM potassium phosphate, 15 mM MgSO4, 4 mM EGTA, and 1 mM DTT) for 20 min at room temperature. Luminescence readings were obtained using a Spectramaxi3X plate reader by mixing 50 μL cell lysate supernatant with 50 μL substrate buffer (lysis buffer with 1 mM ATP, 0.2 mM D-luciferin). Data was normalized to total protein quantified by Bradford assay of the cell lysate.

### Knockdown and knockout cells generation

SMAD2, ATR, ATM, DNAPK, and QKI knockdown JTag cells were generated using pLKO.1 mission shRNAs from Sigma. We generated lentiviral particles from HEK293T by cotransfecting respective shRNA encoding plasmids or control shRNA (shGFP) along with psPAX2, a packaging plasmid and pMD2.G, a VSV glycoprotein encoding plasmid using calcium phosphate method. The medium was replaced after 12-15h post-transfection, and lentiviral particles were collected after 48h. Lentiviral particles containing media was centrifuged at 500g and filtered using syringe filter 0.22 µm. JTAg cells were infected using shRNA particles and selected using puromycin for 1 week. For the generation of TGFBRI-knockout cells, JTag cells were transduced with lentiviral vectors carrying cas9 with guide RNA against TGFBRI. Furthermore, transduced cells were selected for one week with 1 µg/mL puromycin.

### Subcellular fractionation

Subcellular fractionation was done as described previously(Gagnon et al., 2014).For subcellular Fractionation 5 million per condition JTAg cells were collected by spinning down for 500 x g at 4°C for 5 min. Cells were lysed with Hypotonic lysis buffer (HLB) with RNase inhibitor. Cells were kept on ice for 10 mis and then centrifuged at 1000g for 3 min at 4°C. The supernatant was carefully transferred to a new tube with RNA precipitation solution for 2 hours at --20°C and the pellet containing the nucleus was stored in ice. Samples were then centrifuged 18,000 x g at 4°C for 15 min, and the pellet was processed for RNA isolation using TRIzol /chloroform method. The nuclear fraction was washed thrice with ice-cold HLB buffer and centrifuged at 300 x g at 4°C for 2 min.

### Estimation of HIV-1 transcription upon Repsox treatment

JTAg cells were infected by HIV NLBN zsGreen with /without the repsox treatment(10µM). HIV-1 RNA (gag) abundance from cytoplasm as a measure of HIV-1 transcription was quantified at different time points using forward 5’-TTGTACTGAGAGACAGGCT −3’ and Reverse 5’-ACCTGAAGCTCTCTTCTGG-3’. RNA was isolated using TRIzol/Chloroform method and cDNA was synthesized using the random hexamer primer.

### Nuclear Run-on assay

A nuclear run-on assay was performed as described previously(Gagnon et al., 2014). For nuclei isolation, JTAg cells were infected by NL4-3 zsGreen virus with /without the Repsox treatment(10µM) and harvested using ice-cold hypotonic solution (150 mM KCl (Sigma), 4 mM MgOAc (Sigma), and 10 mM Tris-HCl (Sigma), pH 7.4) and were pelleted by centrifugation 300 x g for 5 min. The cells were then lysed in lysis buffer (150 mM KCl, 4 mM MgOAc, 10 mM Tris-HCl, pH 7.4, and 0.5% NP-40). The crude nuclei were mixed with 10 mM ATP, CTP, GTP, BrUTP and incubated at 28 °C for 5 min in the presence of RNase inhibitor. Further the RNA was isolated by TRIzol reagent. The nascent transcripts were then Immunoprecipitated by anti-BrdU antibody (Merck, Cat# B8434) and converted to cDNA for qPCR analysis. qPCR analysis for gag was done using forward 5’-TTGTACTGAGAGACAGGCT −3’ and Reverse 5’-ACCTGAAGCTCTCTTCTGG-3’ primers.

### Immunoblotting

For immunoblotting, samples were lysed using RIPA lysis buffer with 2X PIC and 2mM sodium orthovanadate, 1mM NaF and mixed with Laemmli buffer. Samples were either run on 8% or 15% tricine gels, depending upon protein size. Next, gels were electroblotted on the PVDF membrane (Immobilon-FL, Merck-Millipore). The membrane was blocked using a commercial blocking buffer (BIORAD) for 15 min, followed by primary and secondary antibody incubations for one hour each at room temperature. Post antibody incubation membrane was washed thrice (5 min per wash) using Tween20 containing 1x Tris-Buffered saline (TBST). Details of antibodies are provided in the supplementary table.

### Chromatin Immunoprecipitation (ChIP)

For Chromatin Immunoprecipitation, 10 million JTAg cells per condition were fixed with 1% formaldehyde (Sigma) for 10 minutes to cross-link proteins. The reaction was quenched by adding 0.125 M glycine for 5 minutes at room temperature. The cells were rinsed three times in cold PBS before being suspended in a lysis buffer containing 50 mM HEPES-KOH (pH 7.5), 140 mM NaCl, 1 mM EDTA, 10% glycerol, 0.5% NP-40, 0.25% Triton X-100, with protease inhibitors. The nuclei were centrifuged at 800 × g for 5 minutes at 4 °C and then suspended in lysis buffer containing 10 mM Tris-HCl pH 8.0, 200 mM NaCl, 1 mM EDTA, 0.5 mM EGTA, with protease inhibitors and incubated on ice for 5 minutes. The nuclei were centrifuged at 800g for 5 minutes at 4 °C and then suspended in a buffer consisting of 10 mM Tris-HCl pH 8.0, 200 mM NaCl, 1 mM EDTA, 0.5 mM EGTA, and protease inhibitors and incubated on ice for 10 minutes. The nuclei were gathered and reconstituted in a sonication buffer consisting of 10 mM Tri-HCl (pH 8.0), 100 mM NaCl, 1 mM EDTA, 0.5% EGTA, 0.1% Sodium deoxycholate, and 0.5% N-lauryl sarcosine, along with protease inhibitors.

The DNA was fragmented using a probe sonicator for 15 cycles, with each cycle consisting of 30 seconds of sonication followed by 30 seconds of rest. The samples were centrifuged at 16000g for 10 minutes at a temperature of 4 °C, after treatment with a 1% Triton X-100. For each ChIP cycle, a portion of sonicated DNA was subjected to reverse-crosslinking and then analyzed on a 1% agarose gel to verify the size of the fragments(300-500bp).

Immunoprecipitation was performed on chromatin samples (25 μg) by specific antibodies (anti-smad2/3, anti-Tat, anti-SRSF1, anti-RNAPolII), followed by overnight incubation at 4 °C. Following an incubation period, 30 μl of Protein G beads (Invitrogen) were added and incubated for one hour at a temperature of 4 °C. The beads were sequentially washed for 3 minutes each in low salt (20 mM Tris-HCl pH 8.0, 150 mM NaCl, 2 mM EDTA, 0.1% SDS, 1% Triton X-100), high salt (20 mM Tris-HCl pH 8.0, 500 mM NaCl, 2 mM EDTA, 0.1% SDS, 1% Triton X-100), LiCl buffer (10 mM Tris-HCl pH 8.0, 0.25 M LiCl, 1% NP40, 1% Sodium deoxycholate), and TE buffer. The beads were washed with 150 μl of elution buffer containing 50 mM Tris-HCl pH 8.0, 10 mM EDTA, 1% SDS, and 50 mM NaHCO3. Then, reverse crosslinking was performed with 1 μl of RNaseA (1mg/ml) at a temperature of 37 °C for 30 minutes and subsequently digestion of proteins by adding 1 μl of proteinase K (20mg/ml) and allowing it to react at a temperature of 65°C for 4 hours. The DNA was eluted and used for qPCR analysis.

### Immunofluorescence

HEK293T cells were seeded in 12 well at 1 × 10^5^cells/well with coverslips and allowed to rest overnight. Cells were then transfected with empty vector or Vpr mutant for 20 h. Cells were then fixed in 4% PFA for 20 minutes at room temperature and permeabilized with 0.5% Triton X-100 in PBS for 15 min. Samples were then washed in 1× PBS and incubated with blocking buffer (3% BSA, 0.05% Tween 20, and 0.04 %NaN3 in PBS) for 30 min. Cells were probed with appropriate primary antibodies anti-HA Rabbit (CST), anti-γH2AX (biolegend), and then washed in PBST (0.05% Tween 20 in PBS) thrice for 5mins each and probed with Alexa Fluor-conjugated secondary antibodies. Nuclei were stained with Hoechst 33342(Sigma).

### Intercellular staining of p-smad2/3

For intercellular staining of p-Smad2/3, JTAg cells were challenged with lentiviral particles packaged with HA-Vpr with/without Repsox for 24 hours. The transduced cells were scored with zsgreen positive cells (FITC) where we used p-smad2/3 or respective isotype control in APC channel to ensure the signal was from the transduced cells. Flow cytometry analysis was performed as described earlier(Fujiwara et al., 2023). Briefly, cells were washed with 1XPBS and centrifuged for 5 min at 420g. Cells were fixed by resuspending in 2% paraformaldehyde (PFA) in PBS. Incubate at 37°C for 10 min and centrifuge for 5 min at 420g. Further cells were resuspended cells in 100µL ice-cold methanol. Incubate at 4°C for 60 min in the dark, and cells were washed twice with FACS buffer. We performed intracellular staining by resuspending cells in 100 µL FACS buffer containing phospho-Smad2/3 antibody or isotype antibody as control. Incubate for 60 min at room temperature. Next, cells were washed twice with FACS buffer and secondary antibody Alexa Rabbit 633 was incubated for 30 mins in FACS buffer. Finally, cells were washed thrice with FACS buffer and resuspended in 200µL of FACS buffer for Flow analysis. Data were acquired on BD FACS AriaIII and analyzed in FLOWJO software.

### RNA-Seq Analysis

Analysis of publicly available datasets for TGFB expression in HIV-1 infected patients having variable disease progression (PRJNA824711) and HIV patients with or without ART treatment as compared to healthy individuals (PRJNA984233)(Y. Zhou et al., 2023) was performed. Briefly, for gene expression analysis, we aligned the RNAseq reads to hg38 reference genome using STAR aligner and removed duplicates using samblaster. We generated count matrix using feature counts and then performed differential gene expression analysis using DESeq2 package in R. Graph was plotted DESeq2 normalized counts and performed one-way ANOVA using graphpad prism 9 to determine statistical significance.

### Primary PBMC culture

Primary PBMCs cells were purchased from hiMIdea -HiFiTMHuman Peripheral Blood Mononuclear Cells (Product code CL010) grown and maintained in RPMI-1640 (Gibco) supplemented with 10% FBS. For expanding the CD4+T cells, the cells were either activated with 5 μg/mL phytohemagglutinin (PHA, Sigma-Aldrich) and 50 IU/mL recombinant human IL-2 (Gibco). IL-2 was also supplemented to each culture for the maintenance of the cells. CD3^+^/CD4^+^ T cells were purified by positive selection using magnetic separation with a CD4^+^ isolation kit (Miltenyi Biotec) from PBMCs as per the manufacturer’s protocol. CD4^+^ T cells were by counter-staining with anti-CD4-APC antibody (1:100) using flow cytometry (Miltenyi Biotec), and anti-CD3-FITC labelled antibody. Antibodies were diluted in PBS / 1% BSA / 0.05% NaN3 (PBA).

### ELISA for total TGF-β

PBMCs were treated with NLBN zsGreen virus (with or without 10uM Repsox) for 24 hours in serum free media. Untreated cells were considered as mock samples for the assay. The supernatant was collected from all samples after completion of treatment and centrifuged for 5 mins at 300g. 50ul of the supernatant from all samples was used for the assay. The assay was performed as per the manufacturer’s instructions and a standard curve was generated from the kit components for TGF-β quantification in test samples.

### Cell viability assay

CD4+ cells were treated with Repsox at 10µM. After 24 hours of compound addition, Alamar Blue reagent (HiMedia) was added to each well according to the manufacturer’s protocol, and the plate was incubated at 37°C for 4 h. The absorbance was measured at 570 nm using a SpectraMaxi3X plate reader.

### Conservation of amino acids across Vpr

We obtained alignment of Vpr amino acid sequences derived from patient DNA samples using HIV LANL database (https://www.hiv.lanl.gov/). The alignments were analysed for conservation using Jalview software (73).

### Statistical analysis

Statistical analyses were performed using GraphPad Prism 9.0. Experiments were performed at least 3 times, and data were presented as the means ± standard deviation (SD) from technical replicates. The two-tailed Student’s t-test (unpaired) or one-way ANOVA (multiple comparisons) was used to assess the significance between two or more groups. All reported differences were *p < 0.05, **p < 0.01, ***p < 0.001 and ****p < 0.0001 unless otherwise stated.

## Acknowledgments

A.Cho. received fellowship from IISERB and K.M. and R.D. are supported by fellowships from CSIR. A.Cha. is an EMBO Global Investigator at IISERB. Authors are thankful to M. Pizzato, the NIH AIDS reagent program, and NIBSC for reagents and cell lines. IISERB for infrastructure and all laboratory members, especially V. Bhardwaj, are acknowledged for helpful discussions and technical help.

## Funding and Ethics

This work was supported by the DBT/Wellcome Trust India Alliance Fellowship (IA/I/18/2/504006), EMBO and the Lady Tata Memorial Trust Young Scientist Award to A.Cha. Ethics statement: The Institute Ethics Committee of IISERB approved the study (IISERB/IEC/Certificate/2018-II/04).

## Author contributions

Resources: A.Cha. Data curation: A. Cho., K.M., R.D. and A.Cha. Software: A.Cho., R.D. Formal analysis: A. Cho., K.M., R.D., A.Cha. Validation: V A. Cho., K.M., R.D. Investigation: A. Cho., K.M., R.D. and A.Cha. Methodology: A. Cho., K.M., R.D. Conceptualization, supervision, and funding acquisition: A.Cha.

---

**Supplemetary Figure 1.**
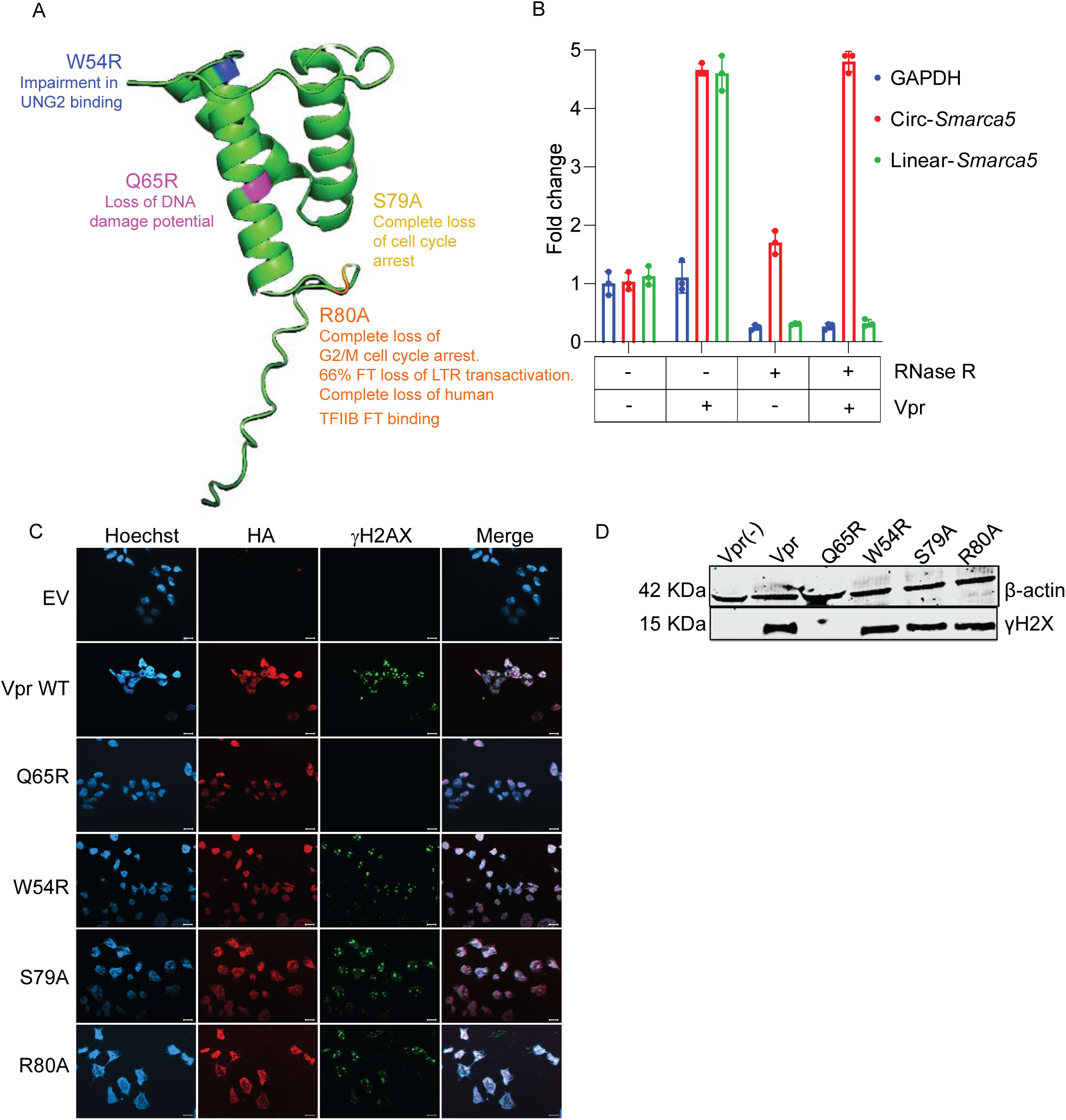
**A.** HIV −1 Vpr (P05928) modelled using Expasy Swissprot representing the mutations studied. **B.** RNAse R treatment was given to RNA isolated from the Vpr (+/-) transduced JTAg cells for 30 mins and levels of circ-smarca5, linear smarca5, GAPDH was assessed by q-RTPCR. All the data was normalized to control RNAseR-samples (n=3±SD). **C.** Immunofluorescence of γH2X with HA Vpr WT and mutants (Scale bar-100μM) **D.** Immunoblot showing the DNA damage marker γH2X with different Vpr mutants with corresponding control as β-actin.

**Supplementary Figure 2.**
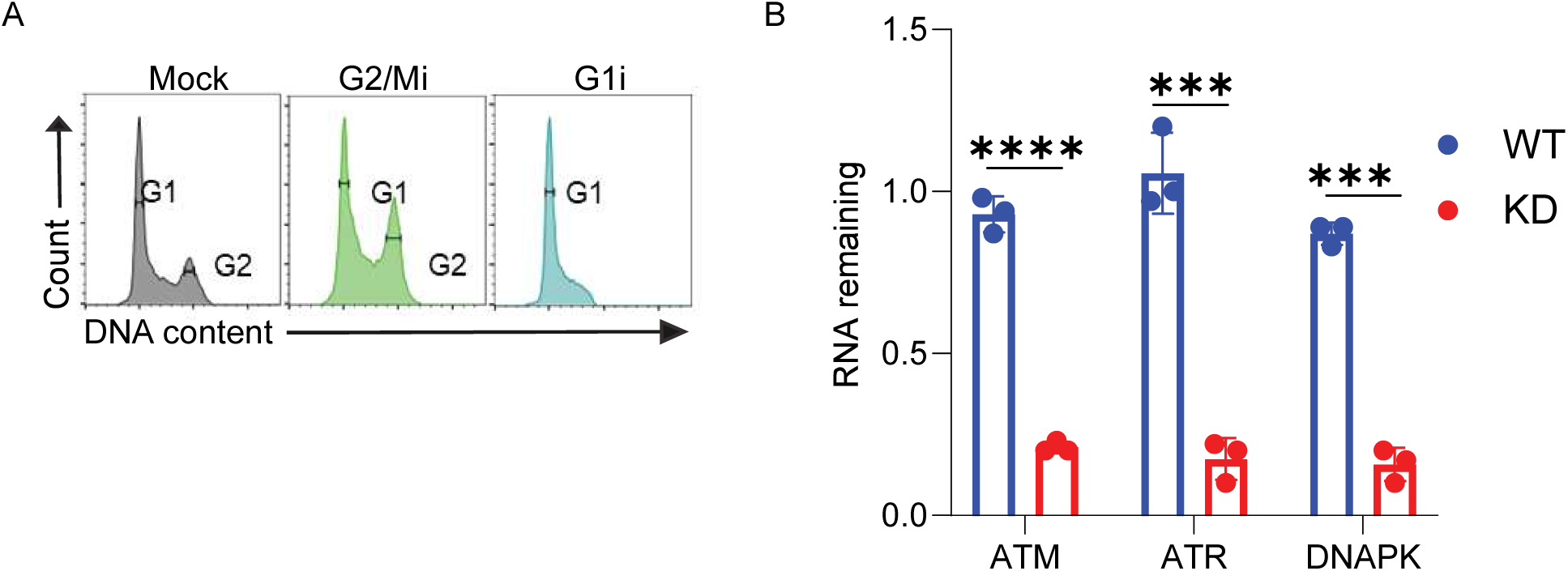
**A.** Flow cytometry analysis of Cell cycle with G2/Mi inhibitor (Apegenin) and G1 inhibitor (CPI 203) and DMSO after 24 hours of treatment in JTAg cells. **B.** Knockdown efficiency of ATM, ATR, DNAPK or JTAg Wild type cells was assessed by qRTPCR. Data were normalized to GAPDH in each conditions (n=3±SD).

**Supplementary Figure 3.**
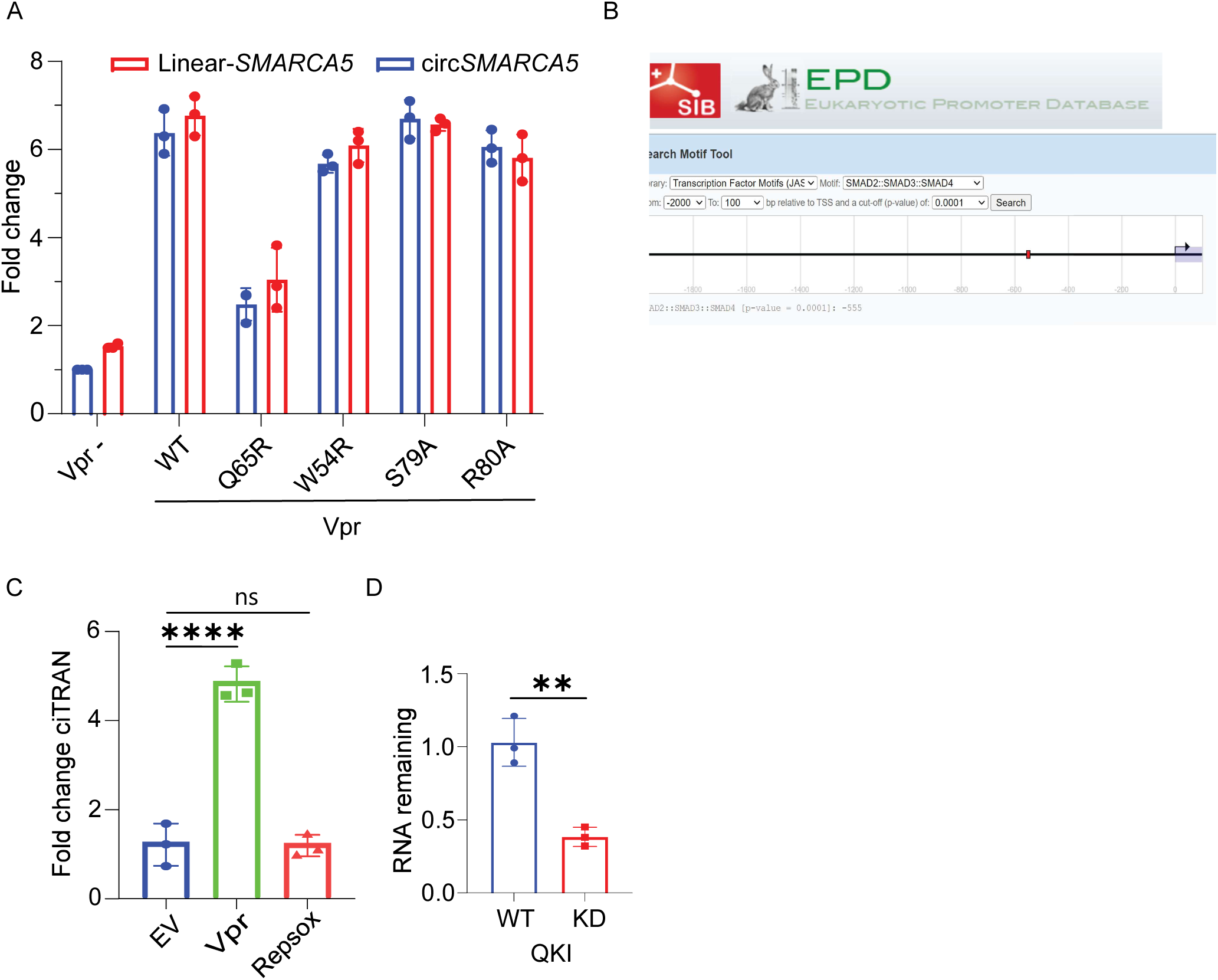
**A.** qRT-PCR of ciTRAN (circ-SMARCA5) and Linear SMARCA5 after transducing JTAg cell with Vpr (-), Vpr (+) LVs (n=3±SD). **B.** A screenshot from the Eukaryotic Promoter database showing binding for the putative binding site of Smad2/3/4. **C.** Effect of Repsox on ciTRAN levels was assessed by qRTPCR in HEK293T cells after 24 hours transfected with either empty vector or Vpr or Repsox (10μM) treatment. **D.** Levels of QKI post-shRNA-mediated knockdown by qRTPCR, data normalized to GAPDH (n=3±SD).

**Supplementary Figure 4.**
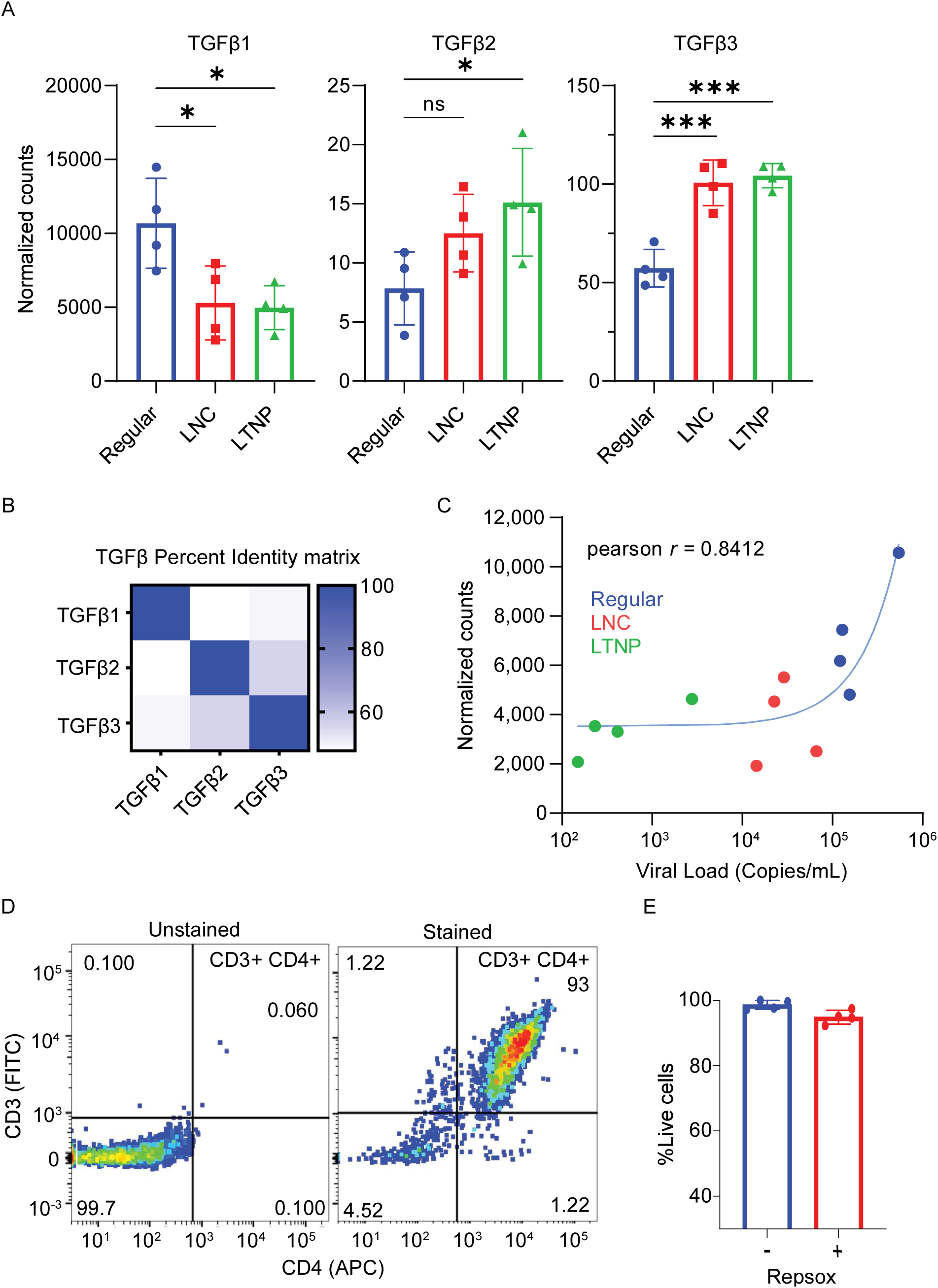
**A.** Levels of TGβ, TGβ2, TGβ3 from the RNAseq data PRJNA824711, from HIV-1 patient samples with variable disease progression. (Regular progressor, LTNP: Long term non progressor, LNC: Long term non-progressor non controller). **B.** TGFB family protein comparison from uniport database as, P01137-TGFB1, P61812-TGFB2, P10600-TGFB3 and aligned using clustal omega to determine their percent identity. **C.** Correlation of TGFβ (normalized count) to Viral load (copies/mL). **D.** Flow cytometry of CD3+/CD4+ cells purified from PBMCs. **E.** Cell viability assay using Alamar blue with /without repsox in human primary CD4+ cells.

**SI Table IA:** List of Plasmids used in the study.

|  |  |  |
| --- | --- | --- |
| 1. pcDNA 3.1 BS (-) | Rosa et al., 2015 | A modified pcDNA derivative with blasticidin selection marker |
| 2. pMD2.G | Addgene# 12259 | Expression of VSV-G |
| 3. psPAX2 | Addgene# 12260 | 2nd generation lentiviral packaging plasmid |
| 4. NLBN zsGreen (HIV-1 zsGreen) (sequence is provided) | Prof. Massimo Pizzato | Modified NL4-3 which is devoid of Env, Nef and ZsGreen is placed in place of Nef |
| 5. pHXB2 env | Rosa et al., 2015 | Expresses Env of HXB2 |
| 6. pCDNA HA | Bhardwaj et al., 2023 | CMV driven expression of HA |
| 7. pCDNA HA Vpr | Bhardwaj et al., 2023 | CMV driven expression of HA-tagged Vpr |
| 8. pCDNA HA Vpr Q65R | Bhardwaj et al., 2023 | CMV driven expression of HA-tagged Vpr mutant Q65R |
| 9. pCDNA HA Vpr W54R | Bhardwaj et al., 2023 | CMV driven expression of HA-tagged Vpr mutant W54R |
| 10. pCDNA HA Vpr R80A | Bhardwaj et al., 2023 | CMV driven expression of HA-tagged Vpr mutant R80A |
| 11. pCDNA HA Vpr S79A | Bhardwaj et al., 2023 | CMV driven expression of HA-tagged Vpr mutant S79A |
| 12. pGL3-Basic | Bhardwaj et al., 2023 | Promoter less vector for measuring the activity of promoter and enhancer sequences with a luciferase assay. |
| 13. pGL3-Basic WT *SMARCA5* Promoter | This Paper | *SMARCA5* Promoter cloned in pGL3 basic vector to check the activity of promoter with luciferase assay |
| 14. pGL3-Basic Mut *SMARCA5* Promoter | This Paper | *Mutant SMARCA5* Promoter cloned in pGL3 basic vector to check the activity of promoter with luciferase assay |
| 15. pLenti CRISre TGFβRI gRNA | Mishra et al., 2022 | For Knockout of TGFβRI |
| 16. pCH040.c/2625 | ARP-11740 | Full-Length Transmitted/Founder (T/F) HIV-1 Infectious Molecular Clones |
| 17. pWITO.c/2474 | ARP-11739 | Full-Length Transmitted/Founder (T/F) HIV-1 Infectious Molecular Clones |
| 18. pSUMA.c/2821 | ARP-11748 | Full-Length Transmitted/Founder (T/F) HIV-1 Infectious Molecular Clones |
| 19. pRHPA.c/2635 | ARP-11744 | Full-Length Transmitted/Founder (T/F) HIV-1 Infectious Molecular Clones |
| 20. pTRJO.c/2851 | ARP-11747 | Full-Length Transmitted/Founder (T/F) HIV-1 Infectious Molecular Clones |
| 21. pREJO.c/2864 | ARP-11746 | Full-Length Transmitted/Founder (T/F) HIV-1 Infectious Molecular Clones |
| 22. pTHRO.c/2626 | ARP-11745 | Full-Length Transmitted/Founder (T/F) HIV-1 Infectious Molecular Clones |
| 23. SBE4-Luc | Addgene# 16495 | SMAD reporter plasmid |
| 24. pGL3 NL4-3 LTR | Bhardwaj et al., 2023 | LTR-Luciferase Reporter |
| 25. pGL3 RHPA LTR | Bhardwaj et al., 2023 | LTR-Luciferase Reporter |
| 26. pGL3 SUMA LTR | Bhardwaj et al., 2023 | LTR-Luciferase Reporter |
| 27. pGL3 TRJO LTR | Bhardwaj et al., 2023 | LTR-Luciferase Reporter |
| 28. pGL3 WITO LTR | Bhardwaj et al., 2023 | LTR-Luciferase Reporter |
| 29. pLenti CRISRe CXCR4 gRNA | Mishra et al., 2022 | For Knockout of CXCR4 |

**SI Table IB:** List of Reagents used in the study.

| SI Table IB: List of Reagents used in the study |  |  |
| --- | --- | --- |
| Reagent | Company | Catalog No. |
| 1. Dulbecco's Modified Eagle Medium (DMEM) | Gibco, USA | 12100046 |
| 2. RPMI 1640 | Gibco, USA | 11875093 |
| 3. Advanced DMEM | Gibco, USA | 12491-015 |
| 4. Advanced RPMI | Gibco, USA | 12633-012 |
| 5. Fetal Bovine Serum (FBS), Certified, Performance tested. Origin: United states | Gibco, USA | 10082-147 |
| 6. Fetal Bovine Serum (FBS), Collected in South America | Capricorn Scientific | FBS12-A |
| 7. L-Glutamax supplement | Gibco, USA | 35050061 |
| 8. Pen-Strep (Penicillin Streptomycin) | Gibco, USA | 15140-122 |
| 9. Trypsin | Gibco | 25200-056 |
| 10. Hoechst 33342 | Sigma Aldrich | 14540 |
| 11. Opti-MEM I | Gibco, USA | 22600-050 |
| 12. PBS | HyClone, USA | SH30256.02 |
| 13. Paraformaldehyde | Sigma Aldrich | F1635-25ML |
| 14. Tris(2-carboxyethyl) phosphine hydrochloride (TCEP) | Sigma Aldrich | 75259 |
| 15. 2-Mercaptoethanol | Sigma | M6250 |
| 16. cOmplete™, EDTA-free Protease Inhibitor Cocktail | Roche | 04693132001 |
| 17. Sodium Orthovanadate | Sigma | S6508 |
| 18. Sodium fluoride | Sigma | 201154 |
| 19. PVDF membrane | Immobilon-FL, Merck-Millipore | IPFL00010 R7DA8781C |
| 20. Tricine | MP Biomedicals | 103112<br>QR15009 |
| 21. Biorad Blocking Buffer | Bio rad | 12010020 |
| 22. Lipofectamine™ 3000 Transfection Reagent | Thermo Fisher Scientific | L3000008 |
| 23. Tween-20® | Sigma Aldrich | P2287-500ML |
| 24. BD Perm/Wash™ Perm/Wash Buffer | BD Biosciences | 554723 |
| 25. Luciferin | Cayman Chemicals | 14682-50MG |
| 26. Dynabeads Protein G | Invitrogen | 10003D |
| 27. SYBR Green-I | Thermo Fisher Scientific | S-7585 |
| 28. TRI Reagent | Sigma | T9424 |
| 29. Revert Aid H minus RT | Thermo Fisher Scientific | K1632 |
| 30. RNase R | Lucigen | RNR07250 |
| 31. α-Amanitin | Sigma Aldrich | B2263 |
| 32. Etoposide | Tocriscreen Plus Micro | 6455 |
| 33. Doxorubicin | Tocriscreen Plus Micro | 6455 |
| 34. Recombinant TGF-β | Gibco | PHG9214 |
| 35. Propidium Iodide | Sigma | 537059 |
| 36. AZ 20 | Tocriscreen Plus Micro | 6455 |
| 37. Apigenin | LOPAC®1280 | LO3300 |
| 38. CPI 203 | Sigma | 6455 |
| 39. Repsox | sigma | R0158-5MG |
| 40. DMSO | Cayman Chemical Company | D8418 |
| 41. KU-55933 | Cayman Chemical Company | 16336 |
| 42. NU 7441 | Cayman Chemical Company | 14881 |
| 43. HiFi™ TM Peripheral Blood Mononuclear cells (H-PBMC),Pooled | HIMEDIA | CL010-10 |
| 44. Gibco™ Human IL-2 Recombinant ProteinL-2 | Gibco | PHC0026 |
| 45. Purified anti-HA.11 Epitope Tag antibody | BioLegend | 901501 |
| 46. Beta-Actin Rabbit Monoclonal Antibody | LI-COR Biosciences | 926-42210 |
| 47. Beta-Actin Mouse Monoclonal Antibody | LI-COR Biosciences | 926-42212 |
| 48. beta Actin Loading Control Monoclonal Antibody (BA3R), DyLight™ 680 | Thermo Fisher Scientific | MA5-15739-D680 |
| 49. IRDye 680RD Goat anti-Mouse IgG antibody | LI-COR Biosciences | 925-68070 |
| 50. IRDye 800CW Goat anti-Rabbit IgG antibody | LI-COR Biosciences | 925-32211 |
| 51. IRDye 680 Donkey anti-Goat IgG antibody | Thermo Fisher Scientific | NC0272752 |
| 52. Anti-HIV-1 p24 Hybridoma | NIH-ARP | 1513 |
| 53. Anti- HIV-1 HXB2 Tat Monoclonal | NIH-ARP | 7377 |
| 54. Rpb1 CTD Mouse Antibody | Cell Signalling Technology | 2629 |
| 55. Human anti-CD4-APC | Miltenyi Biotec | 130-113-812 |
| 56. Human anti-CD3-FITC | Miltenyi Biotec | 130-113-690 |
| 57. Goat TGFβ RI | R&D Systems | AF3025 |
| 58. Rabbit SMAD2/3 | CST | 77397 |
| 59. Rabbit Smad2 | CST | 77397 |
| 60. Rabbit p-Smad2/3 | CST | 77397 |
| 61. Mouse anti-SRSF1 | Thermo Fisher Scientific | 324500 |
| 62. Mouse anti-BrdU | Sigma-Aldrich | B8434 |
| 63. Purified anti-H2A.X Phospho (Ser139) Antibody | Biolegend | 613401 |
| 64. Anti HA rabbit monoclonal antibody(C29F4) | CST | 3724S |
| 65. RNA Clean and Concentrator-5 | Zymo Research | R1014 |
| 66. Nucleospin Gel and PCR clean-up kit | Macherey-Nagel | 740609 |
| 67. NucleoSpin Plasmid Transfection- grade, Mini kit for ultrapure plasmid DNA | Machery-Nagel | 740490 |
| 68. NucleoBond Xtra Midi EF, Midi kit for endotoxin-free plasmid DNA | Machery-Nagel | 740420 |
| 69. NucleoSpin Tissue, Mini kit for DNA from cells and tissue | Machery-Nagel | 740952 |
| 70. Biolegend ELISA TGF-β Kit | Biolegend | 436707 |
| 71. CD4+ T Cell Isolation Kit, human | Miltenyi Biotec | 130-045-101 |
| 72. Nuclease-free water | Thermo Fisher Scientific | AM9937 |

**SI Table IC:** List of Software used in the study.

| SI Table IC: List of Software used in the study |  |  |
| --- | --- | --- |
| 1. GraphPad Prism | GraphPad Software | https://graphpad.com |
| 2. Biorender | NA | https://biorender.com |
| 3. ImageJ |  |  |

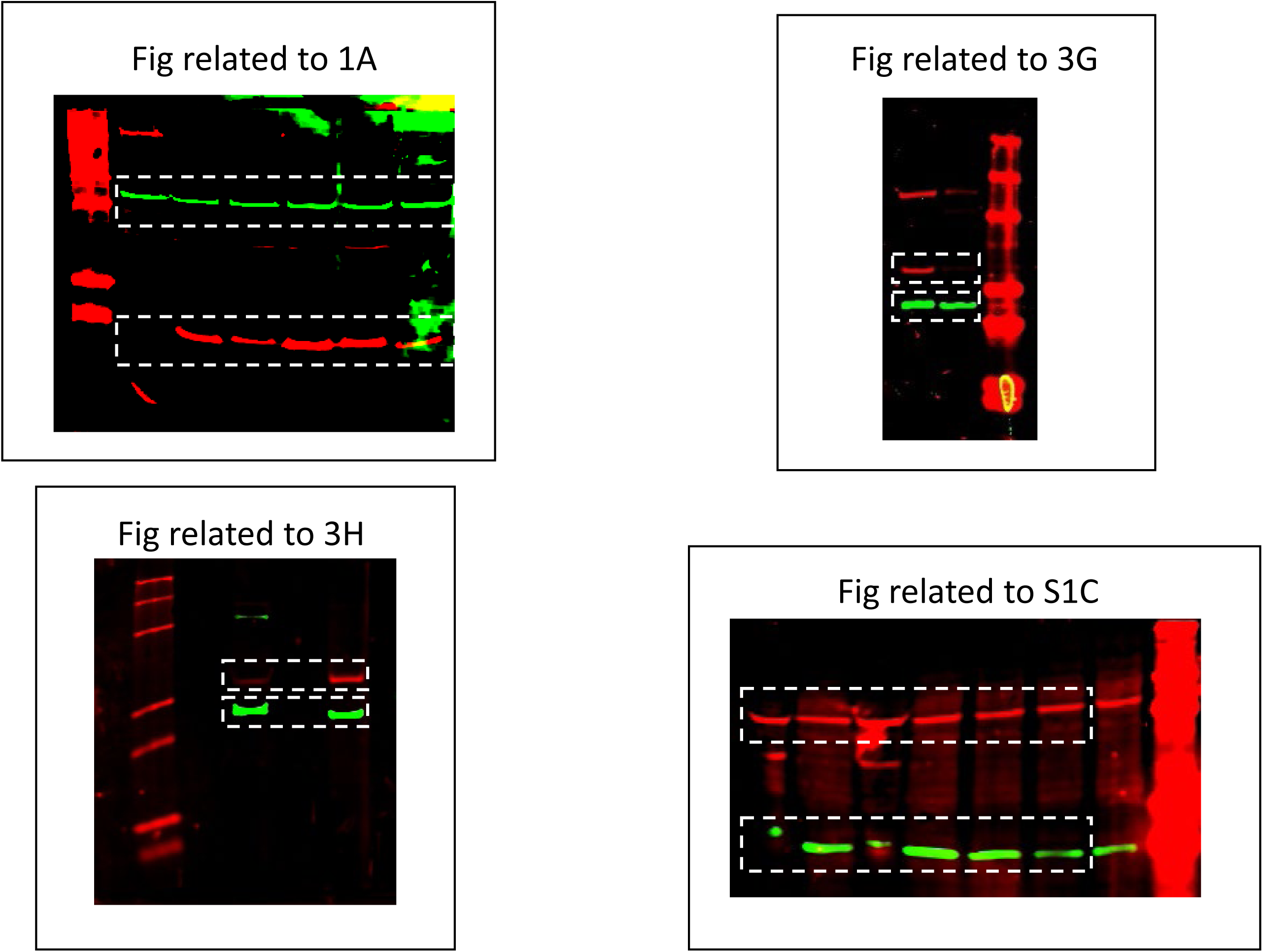

## References

Ali, A., Ng, H. L., Blankson, J. N., Burton, D. R., Buckheit, R. W., Moldt, B., Fulcher, J. A., Javier Ibarrondo, F., Anton, P. A., & Yang, O. O. (2018). Highly attenuated infection with a VPR-deleted molecular clone of human immunodeficiency virus-1. Journal of Infectious Diseases, 218(9), 1447–1452. 10.1093/infdis/jiy346

Barcellos-Hoff, M. H., & Cucinotta, F. A. (2014). New tricks for an old fox: Impact of TGFb on the DNA damage response and genomic stability. Science Signaling, 7(341), 1–6. 10.1126/scisignal.2005474

Belzile, J. P., Duisit, G., Rougeau, N., Mercier, J., Finzi, A., & Cohen, É. A. (2007). HIV-1 Vpr-mediated G2 arrest involves the DDB1-CUL4AVPRBP E3 ubiquitin ligase. PLoS Pathogens, 3(7), 0882– 0893. 10.1371/JOURNAL.PPAT.0030085

Bhardwaj, V., Singh, A., Choudhary, A., Dalavi, R., Ralte, L., Chawngthu, R. L., Kumar, N. S., Vijay, N., & Chande, A. (2023). HIV-1 Vpr induces ciTRAN to prevent transcriptional repression of the provirus. Science Advances, 9(36). 10.1126/SCIADV.ADH9170/SUPPL_FILE/SCIADV.ADH9170_AUXILIARY_DATA_FILES_S1_AND_S2.ZIP

Bhardwaj, V., Singh, A., Dalavi, R., Ralte, L., Chawngthu, R. L., Kumar, N. S., Vijay, N., & Chande, A. (2022). HIV-1 Vpr induces ciTRAN to prevent transcriptional silencing of the provirus. BioRxiv, 2022.11.04.515166. https://www.biorxiv.org/content/10.1101/2022.11.04.515166v1 https://www.biorxiv.org/content/10.1101/2022.11.04.515166v1.abstract

Chen, L. L. (2020). The expanding regulatory mechanisms and cellular functions of circular RNAs. Nature Reviews Molecular Cell Biology, 21(8), 475–490. 10.1038/S41580-020-0243-Y

Chinnapaiyan, S., Dutta, R. K., Nair, M., Chand, H. S., Rahman, I., & Unwalla, H. J. (2019). TGF-β1 increases viral burden and promotes HIV-1 latency in primary differentiated human bronchial epithelial cells. Scientific Reports, 9(1), 1–10. 10.1038/s41598-019-49056-6

Choudhary, A., Madbhagat, P., Sreepadmanabh, M., Bhardwaj, V., & Chande, A. (2021). Circular RNA as an Additional Player in the Conflicts Between the Host and the Virus. Frontiers in Immunology, 12. 10.3389/FIMMU.2021.602006

Cohen, E. A., Dehni, G., Sodroski, J. G., & Haseltine, W. A. (1990). Human immunodeficiency virus vpr product is a virion-associated regulatory protein. Journal of Virology, 64(6), 3097–3099. 10.1128/jvi.64.6.3097-3099.1990

Cohen, E. A., Terwilliger, E. F., Jalinoos, Y., Proulx, J., Sodroski, J. G., & Haseltine, W. A. (1990). Identification of HIV-1 vpr product and function. Journal of Acquired Immune Deficiency Syndromes, 3(1), 11–18.

Dickinson, M., Kliszczak, A. E., Giannoulatou, E., Peppa, D., Pellegrino, P., Williams, I., Drakesmith, H., & Borrow, P. (2020). Dynamics of Transforming Growth Factor (TGF)-β Superfamily Cytokine Induction During HIV-1 Infection Are Distinct From Other Innate Cytokines. Frontiers in Immunology, 11(November), 1–16. 10.3389/fimmu.2020.596841

Dong, R., Zhang, X. O., Zhang, Y., Ma, X. K., Chen, L. L., & Yang, L. (2016). CircRNA-derived pseudogenes. Cell Research, 26(6), 747–750. 10.1038/cr.2016.42

Edgar, R. C., Taylor, J., Lin, V., Altman, T., Barbera, P., Meleshko, D., Lohr, D., Novakovsky, G., Buchfink, B., Al-Shayeb, B., Banfield, J. F., de la Peña, M., Korobeynikov, A., Chikhi, R., & Babaian, A. (2022). Petabase-scale sequence alignment catalyses viral discovery. Nature, 602(7895), 142–147. 10.1038/s41586-021-04332-2

Forget, J., Yao, X. J., Mercier, J., & Cohen, É. A. (1998). Human immunodeficiency virus type 1 Vpr protein transactivation function: Mechanism and identification of domains involved. Journal of Molecular Biology, 284(4), 915–923. 10.1006/jmbi.1998.2206

Fu, S., Phan, A. T., Mao, D., Wang, X., Gao, G., Goff, S. P., & Zhu, Y. (2022). HIV-1 exploits the Fanconi anemia pathway for viral DNA integration. 39(8). 10.1016/j.celrep.2022.110840.HIV-1

Fujiwara, M., Amrendra, K., & Alkeiver, S. (2023). Protocol Protocol for analyzing transforming growth factor b signaling in dextran-sulfate-sodium-induced colitic mice using flow cytometry and western blotting Protocol for analyzing transforming growth factor b signaling in dextran-sulfate-sodium-induce. STAR Protocols, 4(2), 102249. 10.1016/j.xpro.2023.102249

Gagnon, K. T., Li, L., Janowski, B. A., & Corey, D. R. (2014). Analysis of nuclear RNA interference in human cells by subcellular fractionation and Argonaute loading. Nature Protocols, 9(9), 2045– 2060. 10.1038/nprot.2014.135

Goh, W. C., Rogel, M. E., Matthew Kinsey, C., Michael, S. F., Fultz, P. N., Nowak, M. A., Hahn, B. H., & Emerman, M. (1998). HIV-1 Vpr increases viral expression by manipulation of the cell cycle: A mechanism for selection of Vpr in vivo. Nature Medicine, 4(1), 65–71. 10.1038/NM0198-065

Gopalakrishnan, R. M., Aid, M., Mercado, N. B., Davis, C., Malik, S., Geiger, E., Varner, V., Jones, R., Bosinger, S. E., Piedra-Mora, C., Martinot, A. J., Barouch, D. H., Keith Reeves, R., & Sabrina Tan, C. (2021). Increased IL-6 expression precedes reliable viral detection in the rhesus macaque brain during acute SIV infection. JCI Insight, 6(20). 10.1172/jci.insight.152013

Guenzel, C. A., Hérate, C., & Benichou, S. (2014). HIV-1 Vpr-a still “enigmatic multitasker.” Frontiers in Microbiology, 5(MAR), 1–13. 10.3389/fmicb.2014.00127

Guenzel, C. A., Hérate, C., Le Rouzic, E., Maidou-Peindara, P., Sadler, H. A., Rouyez, M.-C., Mansky, L. M., & Benichou, S. (2012). Recruitment of the Nuclear Form of Uracil DNA Glycosylase into Virus Particles Participates in the Full Infectivity of HIV-1. Journal of Virology, 86(5), 2533–2544. 10.1128/jvi.05163-11

Gyori, B. M., Venkatachalam, G., Thiagarajan, P. S., Hsu, D., & Clement, M.-V. (2014). OpenComet: An automated tool for comet assay image analysis. Redox Biology, 2, 457–465. 10.1016/j.redox.2013.12.020

Hansen, T. B., Jensen, T. I., Clausen, B. H., Bramsen, J. B., Finsen, B., Damgaard, C. K., & Kjems, J. (2013). Natural RNA circles function as efficient microRNA sponges. Nature, 495(7441), 384–388. 10.1038/nature11993

Harman, A. N., Nasr, N., Feetham, A., Galoyan, A., Alshehri, A. A., Rambukwelle, D., Botting, R. A., Hiener, B. M., Diefenbach, E., Diefenbach, R. J., Kim, M., Mansell, A., & Cunningham, A. L. (2015). HIV Blocks Interferon Induction in Human Dendritic Cells and Macrophages by Dysregulation of TBK1. J Virol., 89(13), 6575–6584. 10.1128/jvi.00889-15

He, J., Choe, S., Walker, R., Di Marzio, P., Morgan, D. O., & Landau, N. R. (1995). Human immunodeficiency virus type 1 viral protein R (Vpr) arrests cells in the G2 phase of the cell cycle by inhibiting p34cdc2 activity. Journal of Virology, 69(11), 6705–6711. 10.1128/JVI.69.11.6705-6711.1995

Hrimech, M., Yao, X.-J., Bachand, F., Rougeau, N., & Cohen, É. A. (1999). Human Immunodeficiency Virus Type 1 (HIV-1) Vpr Functions as an Immediate-Early Protein during HIV-1 Infection. Journal of Virology, 73(5), 4101–4109. 10.1128/jvi.73.5.4101-4109.1999

Iijima, K., Kobayashi, J., & Ishizaka, Y. (2018). Structural alteration of DNA induced by viral protein R of HIV-1 triggers the DNA damage response. Retrovirology, 15(1), 1–25. 10.1186/s12977-018-0391-8

Jeck, W. R., Sorrentino, J. A., Wang, K., Slevin, M. K., Burd, C. E., Liu, J., Marzluff, W. F., & Sharpless, N. E. (2013). Circular RNAs are abundant, conserved, and associated with ALU repeats. RNA, 19(2), 141–157. 10.1261/rna.035667.112

Kekow, J., Wachsman, W., Allen McCutchan, J., Gross, W. L., Zachariah, M., Carson, D. A., & Lotz, M. (1991). Transforming growth factor-β and suppression of humoral immune responses in HIV infection. Journal of Clinical Investigation, 87(3), 1010–1016. 10.1172/jci115059

Kekow, J., Wachsman, W., McCutchan, J. A., Cronin, M., Carson, D. A., & Lotz, M. (1990). Transforming growth factor β and noncytopathic mechanisms of immunodeficiency in human immunodeficiency virus infection. Proceedings of the National Academy of Sciences of the United States of America, 87(21), 8321–8325. 10.1073/pnas.87.21.8321

Khan, A., Fornes, O., Stigliani, A., Gheorghe, M., Lee, R. Van Der, Bessy, A., Ch, J., Tan, G., Baranasic, D., Arenillas, D. J., Sandelin, A., Vandepoele, K., & Lenhard, B. (2018). JASPAR 2018: update of the open-access database of transcription factor binding profiles and its web framework. 46(November 2017), 260–266. 10.1093/nar/gkx1126

Kim, J., Bose, D., Araínga, M., Haque, M. R., Fennessey, C. M., Caddell, R. A., Thomas, Y., Ferrell, D. E., Ali, S., Grody, E., Goyal, Y., Cicala, C., Arthos, J., Keele, B. F., Vaccari, M., Lorenzo-Redondo, R., Hope, T. J., Villinger, F., & Martinelli, E. (2024). TGF-β blockade drives a transitional effector phenotype in T cells reversing SIV latency and decreasing SIV reservoirs in vivo. Nature Communications, 15(1), 1–17. 10.1038/s41467-024-45555-x

Lasda, E., & Parker, R. (2014). Circular RNAs: Diversity of form and function. In RNA (Vol. 20, Issue 12, pp. 1829–1842). Cold Spring Harbor Laboratory Press. 10.1261/rna.047126.114

Lazdins, J. K., Klimkait, T., Woods-Cook, K., Walker, M., Alteri, E., Cox, D., Cerletti, N., Shipman, R., Bilbe, G., & Mcmaster, G. (1992). The Replicative Restriction of Lymphocytotropic Isolates of HIV-1 in Macrophages Is Overcome by TGF-β. AIDS Research and Human Retroviruses, 8(4), 505–511. 10.1089/aid.1992.8.505

Le Rouzic, E., Belaïdouni, N., Estrabaud, E., Morel, M., Rain, J. C., Transy, C., & Margottin-Goguet, F. (2007). HIV1 Vpr arrests the cell cycle by recruiting DCAF1/VprBP, a receptor of the Cul4-DDB1 ubiquitin ligase. Cell Cycle, 6(2), 182–188. 10.4161/CC.6.2.3732

Lee, B. D., Neri, U., Roux, S., Wolf, Y. I., Camargo, A. P., Krupovic, M., Simmonds, P., Kyrpides, N., Gophna, U., Dolja, V. V., & Koonin, E. V. (2023). Mining metatranscriptomes reveals a vast world of viroid-like circular RNAs. Cell, 186(3), 646–661.e4. 10.1016/j.cell.2022.12.039

Li, D., Lopez, A., Sandoval, C., Doyle, R. N., & Fregoso, O. I. (2020). HIV VPR modulates the host DNA damage response at two independent steps to damage dna and repress double-strand dna break repair. MBio, 11(4), 1–18. 10.1128/mBio.00940-20

Li, Y., Liu, Y., Chiang, Y. J., Huang, F., Li, Y., Li, X., Ning, Y., Zhang, W., Deng, H., & Chen, Y. G. (2019). DNA Damage Activates TGF-β Signaling via ATM-c-Cbl-Mediated Stabilization of the Type II Receptor TβRII. Cell Reports, 28(3), 735–745.e4. 10.1016/j.celrep.2019.06.045

Liu, Q., Palomero, L., Moore, J., Guix, I., Espín, R., Aytés, A., Mao, J. H., Paulovich, A. G., Whiteaker, J. R., Ivey, R. G., Iliakis, G., Luo, D., Chalmers, A. J., Murnane, J., Pujana, M. A., & Barcellos-Hoff, M. H. (2021). Loss of TGFB signaling increases alternative end-joining DNA repair that sensitizes to genotoxic therapies across cancer types. Science Translational Medicine, 13(580). 10.1126/scitranslmed.abc4465

Lotz, M., & Seth, P. (1993). TGFβ and HIV Infection. Annals of the New York Academy of Sciences, 685(1), 501–511. 10.1111/j.1749-6632.1993.tb35912.x

Maiuri, T., Mocle, A. J., Hung, C. L., Xia, J., van Roon-Mom, W. M. C., & Truant, R. (2017). Huntingtin is a scaffolding protein in the ATM oxidative DNA damage response complex. Human Molecular Genetics, 26(2), 395–406. 10.1093/hmg/ddw395

Malim, M. H., & Emerman, M. (2008). HIV-1 Accessory Proteins-Ensuring Viral Survival in a Hostile Environment. Cell Host and Microbe, 3(6), 388–398. 10.1016/j.chom.2008.04.008

Maudet, C., Bertrand, M., Le Rouzic, E., Lahouassa, H., Ayinde, D., Nisole, S., Goujon, C., Cimarelli, A., Margottin-Goguet, F., & Transy, C. (2011). Molecular insight into how HIV-1 Vpr protein impairs cell growth through two genetically distinct pathways. Journal of Biological Chemistry, 286(27), 23742–23752. 10.1074/jbc.M111.220780

Memczak, S., Jens, M., Elefsinioti, A., Torti, F., Krueger, J., Rybak, A., Maier, L., Mackowiak, S. D., Gregersen, L. H., Munschauer, M., Loewer, A., Ziebold, U., Landthaler, M., Kocks, C., Le Noble, F., & Rajewsky, N. (2013). Circular RNAs are a large class of animal RNAs with regulatory potency. Nature, 495(7441), 333–338. 10.1038/nature11928

Meylan, P., Ambrosini, G., & Groux, R. (2020). EPD in 2020: enhanced data visualization and extension to ncRNA promoters. 48(November 2019), 65–69. 10.1093/nar/gkz1014

Mishra, T., Bhardwaj, V., Ahuja, N., Gadgil, P., Ramdas, P., Shukla, S., & Chande, A. (2022). Improved loss-of-function CRISPR-Cas9 genome editing in human cells concomitant with inhibition of TGF-β signaling. Molecular Therapy - Nucleic Acids, 28(June), 202–218. 10.1016/j.omtn.2022.03.003

Moustakas, A., Pardali, K., Gaal, A., & Heldin, C.-H. (2002). Mechanisms of TGF-β signaling in regulation of cell growth and differentiation. Immunology Letters, 82(1), 85–91. 10.1016/S0165-2478(02)00023-8

Nakao, A., Imamura, T., Souchelnytskyi, S., Kawabata, M., Ishisaki, A., Oeda, E., Tamaki, K., Hanai, J. I., Heldin, C. H., Miyazono, K., & Ten Dijke, P. (1997). TGF-β receptor-mediated signalling through Smad2, Smad3 and Smad4. EMBO Journal, 16(17), 5353–5362. 10.1093/emboj/16.17.5353

O’Reilly, S., Ciechomska, M., Cant, R., & Van Laar, J. M. (2014). Interleukin-6 (IL-6) trans signaling drives a STAT3-dependent pathway that leads to hyperactive transforming growth factor-β (TGF-β) signaling promoting SMAD3 activation and fibrosis via gremlin protein. Journal of Biological Chemistry, 289(14), 9952–9960. 10.1074/jbc.M113.545822

Paz, S., Krainer, A. R., & Caputi, M. (2014). HIV-1 transcription is regulated by splicing factor SRSF1. Nucleic Acids Research, 42(22), 13812–13823. 10.1093/NAR/GKU1170

Pizzato, M., Erlwein, O., Bonsall, D., Kaye, S., Muir, D., & McClure, M. O. (2009). A one-step SYBR Green I-based product-enhanced reverse transcriptase assay for the quantitation of retroviruses in cell culture supernatants. Journal of Virological Methods, 156(1–2), 1–7. 10.1016/J.JVIROMET.2008.10.012

Pizzato, M., Helander, A., Popova, E., Calistri, A., Zamborlini, A., Palù, G., & Göttlinger, H. G. (2007). Dynamin 2 is required for the enhancement of HIV-1 infectivity by Nef. Proceedings of the National Academy of Sciences of the United States of America, 104(16), 6812–6817. 10.1073/PNAS.0607622104

Rosa, A., Chande, A., Ziglio, S., De Sanctis, V., Bertorelli, R., Goh, S. L., McCauley, S. M., Nowosielska, A., Antonarakis, S. E., Luban, J., Santoni, F. A., & Pizzato, M. (2015). HIV-1 Nef promotes infection by excluding SERINC5 from virion incorporation. Nature, 526(7572), 212–217. 10.1038/NATURE15399

Roth, W. K. (1993). TGFβ and FGF-like growth factors involved in the pathogenesis of AIDS-associated Kaposi’s sarcoma. Research in Virology, 144, 105–109. 10.1016/S0923-2516(06)80019-1

Salzman, J., Gawad, C., Wang, P. L., Lacayo, N., & Brown, P. O. (2012). Circular RNAs are the predominant transcript isoform from hundreds of human genes in diverse cell types. PLoS ONE, 7(2). 10.1371/journal.pone.0030733

Schröfelbauer, B., Hakata, Y., & Landau, N. R. (2007). HIV-1 Vpr function is mediated by interaction with the damage-specific DNA-binding protein DDB1. Proceedings of the National Academy of Sciences of the United States of America, 104(10), 4130–4135. 10.1073/pnas.0610167104

Starke, S., Jost, I., Rossbach, O., Schneider, T., Schreiner, S., Hung, L. H., & Bindereif, A. (2015). Exon circularization requires canonical splice signals. Cell Reports, 10(1), 103–111. 10.1016/j.celrep.2014.12.002

Subbramanian, R. A., Kessous-Elbaz, A., Lodge, R., Forget, J., Yao, X. J., Bergeron, D., & Cohen, E. A. (1998). Human immunodeficiency virus type 1 Vpr is a positive regulator of viral transcription and infectivity in primary human macrophages. Journal of Experimental Medicine, 187(7), 1103–1111. 10.1084/JEM.187.7.1103

Tachiwana, H., Shimura, M., Nakai-Murakami, C., Tokunaga, K., Takizawa, Y., Sata, T., Kurumizaka, H., & Ishizaka, Y. (2006). HIV-1 Vpr induces DNA double-strand breaks. Cancer Research, 66(2), 627–631. 10.1158/0008-5472.CAN-05-3144

Tagawa, T., Gao, S., Koparde, V. N., Gonzalez, M., Spouge, J. L., & Serquiña, A. P. (2018). Discovery of Kaposi’s sarcoma herpesvirus-encoded circular RNAs and a human antiviral circular RNA. 115(50), 12805–12810. 10.1073/pnas.1816183115

Ungerleider, N. A., Jain, V., Wang, Y., Maness, N. J., Blair, R. V., Alvarez, X., Midkiff, C., Kolson, D., Bai, S., Roberts, C., Moss, W. N., Wang, X., Serfecz, J., Seddon, M., Lehman, T., Ma, T., Dong, Y., Renne, R., Tibbetts, S. A., & Flemington, E. K. (2019). Comparative Analysis of Gammaherpesvirus Circular RNA Repertoires: Conserved and Unique Viral Circular RNAs. Journal of Virology, 93(6). 10.1128/jvi.01952-18

Wang, P. L., Bao, Y., Yee, M., Barrett, S. P., Hogan, G. J., Dinneny, R., Brown, P. O., & Salzman, J. (2014). Circular RNA Is Expressed across the Eukaryotic Tree of Life. 9(3). 10.1371/journal.pone.0090859

Wang, W., Sun, L., Huang, M. T., Quan, Y., Jiang, T., Miao, Z., & Zhang, Q. (2023). Regulatory circular RNAs in viral diseases: applications in diagnosis and therapy. RNA Biology, 20(1), 847–858. 10.1080/15476286.2023.2272118

Wiercińska-Drapalo, A., Flisiak, R., Jaroszewicz, J., & Prokopowicz, D. (2004). Increased Plasma Transforming Growth Factor-β1 Is Associated with Disease Progression in HIV-1-Infected Patients. Viral Immunology, 17(1), 109–113. 10.1089/088282404322875502

Xiao, L. Z., Topley, N., Ito, T., & Phillips, A. (2005). Interleukin-6 regulation of transforming growth factor (TGF)-β receptor compartmentalization and turnover enhances TGF-β1 signaling. Journal of Biological Chemistry, 280(13), 12239–12245. 10.1074/jbc.M413284200

Xu, X., Zhang, J., Tian, Y., Gao, Y., Dong, X., Chen, W., Yuan, X., Yin, W., Xu, J., Chen, K., He, C., & Wei, L. (2020). CircRNA inhibits DNA damage repair by interacting with host gene. Molecular Cancer, 19(1), 1–20. 10.1186/s12943-020-01246-x

Yim, L. Y., Lam, K. S., Luk, T.-Y., Mo, Y., Lu, X., Wang, J., Cheung, K.-W., Lui, G. C. Y., Chan, D. P. C., Wong, B. C. K., Lau, T. T.-K., Ngan, C. B., Zhou, D., Wong, Y. C., Tan, Z., Liu, L., Wu, H., Zhang, T., Lee, S. S., & Chen, Z. (2023). Transforming Growth Factor β Signaling Promotes HIV-1 Infection in Activated and Resting Memory CD4+ T Cells. Journal of Virology. 10.1128/JVI.00270-23

Zhang, M., Zhang, Y. Y., Chen, Y., Wang, J., Wang, Q., & Lu, H. (2021). TGF-β Signaling and Resistance to Cancer Therapy. Frontiers in Cell and Developmental Biology, 9(November), 1–15. 10.3389/fcell.2021.786728

Zhang, X., Wang, H., Zhang, Y., Lu, X., Chen, L., & Yang, L. (2014). Complementary Sequence-Mediated Exon Circularization. Cell, 159(1), 134–147. 10.1016/j.cell.2014.09.001

Zhou, D., Munster, A., & Winchurch, R. A. (1991). Pathologic concentrations of interleukin 6 inhibit T cell responses via induction of activation of TGF-β. The FASEB Journal, 5(11), 2582–2585. 10.1096/fasebj.5.11.1868982

Zhou, Y., Huang, Y., Chen, X., Chen, T., Hu, W., Hou, W., Zhang, Q., & Xiong, Y. (2023). Transcriptomic study reveals changes of lncRNAs in PBMCs from HIV-1 patients before and after ART. Scientific Reports, 13(1), 1–12. 10.1038/s41598-023-49595-z

Zimmerman, E. S., Sherman, M. P., Blackett, J. L., Neidleman, J. A., Kreis, C., Mundt, P., Williams, S. A., Warmerdam, M., Kahn, J., Hecht, F. M., Grant, R. M., de Noronha, C. M. C., Weyrich, A. S., Greene, W. C., & Planelles, V. (2006). Human Immunodeficiency Virus Type 1 Vpr Induces DNA Replication Stress InVitro and In Vivo. Journal of Virology, 80(21), 10407–10418. 10.1128/jvi.01212-06

